# Mitochondrial rewiring supports survival of triple negative breast cancer cells after ionizing radiation

**DOI:** 10.64898/2026.09.03.748963

**Authors:** Steven W. Wall, Karen Wang, Allison Greer, Mokryun L. Baek, Audra Lane, Mariah J. Berner, Jonathan T. Lei, Ángel A. Garcés, Sofia Khoury, Emily G. Caggiano, Matthew D. Meyer, Sarah Latka, Hugo Villanueva, Emil Schüler, Cullen M. Taniguchi, Rutulkumar Patel, Gloria V. Echeverria

## Abstract

Triple negative breast cancer (TNBC) is an aggressive disease with limited therapeutic options. Conventional treatments include neoadjuvant chemo-immunotherapy followed by surgical resection and may include further adjuvant immunotherapy and/or radiotherapy of the tumor bed and lymph nodes. Nonetheless, TNBC patients with residual disease have rapid metastatic recurrence. While the roles of metabolic and mitochondrial adaptations in chemotherapeutic resistance have been the subject of many studies, their importance in the context of ionizing radiation (IR) therapy remains poorly understood. We established longitudinal in vitro models of post-IR human TNBC, characterized by cellular regression to a residual phenotypic state, then eventual cell repopulation. This was accompanied by plastic adoption of unique metabolic, proteomic, and morphologic features that largely reverted when cells regrew. Following IR, residual cells exhibited extensive mitochondrial rewiring, including elevated mitochondrial content, oxidative phosphorylation (oxphos) rates, cristae structures, and metabolite levels. Concomitantly, levels of the short protein isoform of the mitochondrial inner membrane protein optic atrophy 1 (OPA1) were significantly elevated in residual cells, and OPA1 knockout ablated mitochondrial adaptations induced by IR. OPA1 genetic or pharmacologic perturbations led to improved cellular responses to IR. Metabolomic and proteomic analyses of radio-residual cells uncovered a coordinated program of antioxidant and redox capacity elevation with mitochondrial metabolism, which was corroborated by analyses of external datasets. Together, these findings provide evidence that TNBC cells surviving radiotherapy adopt an OPA1-dependent program of mitochondrial reorganization that supports their survival and regrowth, thereby positioning OPA1 as a therapeutic dependency that could improve radiotherapy efficacy in TNBC.

## INTRODUCTION

Triple negative breast cancer (TNBC) is a subtype of breast cancer characterized by its lack of the progesterone receptor, the estrogen receptor, and the lack of overexpression of human epidermal growth factor receptor 2^1^. TNBC, having limited targeted therapy options, is routinely managed by systemic neoadjuvant chemo-immunotherapy, with recent addition of chemotherapy-payload antibody-drug conjugates in the metastatic setting^2–4^. Radiotherapy (RT) is a cornerstone of locoregional management in TNBC: it is delivered after breast-conserving surgery in nearly all patients and after mastectomy in those with node-positive disease, large primary tumors, or residual disease following neoadjuvant chemotherapy^5^. Regional nodal irradiation further reduces locoregional recurrence in node-positive patients, and because TNBC carries a disproportionately high recurrence risk relative to other breast cancer subtypes, some expert panels now favor post-mastectomy radiotherapy even after a pathologic complete response^6^. Despite these intensive therapeutic regimens, those ∼40-45% of TNBC patients with residual disease at the time of surgery have high incidence of rapid metastatic recurrence ^7,8^. Unfortunately, the five-year survival rate of metastasized TNBC is only 15%^9^. Thus, understanding the mechanisms that drive treatment resistance in TNBC is critical to developing more effective treatment strategies for standard-of-care-refractory disease.

Mitochondrial adaptations are crucial in facilitating chemotherapy resistance in TNBC ^10,11^. We previously demonstrated TNBC cells surviving conventional DNA-damaging chemotherapy treatments exhibited mitochondrial adaptations such as increased oxidative phosphorylation (oxphos), mitochondrial DNA (mtDNA) content, and mitochondrial fusion driven by changes in optic atrophy 1 (OPA1)^12^. The importance of mitochondria in chemoresistant TNBC has been further affirmed in baseline and longitudinal analyses of pre-and post-chemotherapy human biopsies^13^. Furthermore, targeting cells with the OPA1 inhibitor, MYLS22^14^, or with an oxphos Complex I inhibitor, IACS-010759^15^, significantly improved chemotherapeutic efficacy^10,11^. The nature, functional relevance, and therapeutic tractability of mitochondrial adaptations in TNBCs that are refractory to conventional radiotherapy remain poorly understood.

In this study, we establish *in vitro* models to longitudinally study radio-refractory TNBC and discover pronounced, plastic adaptation of mitochondrial form and function accompanied by extensive metabolic rewiring. Using this model, we investigated mitochondrial adaptations in ‘radio residual’ cells and found OPA1 plays an integral role in promoting the mitochondrial state and regrowth of residual cells. Our findings, corroborated by external TNBC datasets, support a role for OPA1 in post-IR survival and warrant further investigation into leveraging OPA1 and other metabolic inhibitors to abate TNBC recurrence after radiotherapy. This study mechanistically implicates OPA1, cristae remodeling, and metabolic reprogramming as crucial for survival of radio-residual TNBC cells.

## METHODS

### Cell culture and authentication

MDA-MB-231 cells were purchased from American Type Culture Collection (ATCC), and SUM159pt cells were purchased from BioIVT. All cell lines were cultured at 37°C with 5% CO_2_ and 95% relative humidity. MDA-MB-231 cells were cultured in RPMI-1640 (Gibco, 11879020) supplemented with 10% heat-inactivated fetal bovine serum (R&D, S11550), 5 mM glucose (Gibco, A2494001), and 1X antibiotic-antimycotic solution (Corning, 30-004-CI) (231 growth medium). SUM159pt cells were cultured in RPMI-1640 (Gibco, 11879020) supplemented with 5% fetal bovine serum (R&D, S11550), 5 mM glucose (Gibco, A2494001), 1X antibiotic-antimycotic solution (Corning, 30-004-CI), 5 µg/mL insulin (Sigma-Aldrich, I9278) and 1 µg/mL hydrocortisone (Sigma-Aldrich, H4001) (159 growth medium).

Cell lines were tested for mycoplasma contamination each quarter by PCR with the Universal Mycoplasma detection kit (ATCC, 30-1012K). All cells remained mycoplasma free during these studies.

Short-tandem repeat DNA fingerprinting was performed on DNA extracted from cell lines by the Cytogenetics and Cell Authentication Core at M.D. Anderson Cancer Center. The Promega 16 High Sensitivity STR Kit (Catalog # DC2100) was used for fingerprinting analysis, and profiles were compared to online search databases (DSMZ/ ATCC/ JCRB/ RIKEN).

### Gene silencing

We deleted two small coding exons of OPA1 in MDA-MB-231 cells using CRISPR/Cas9, and the deletion resulted in a frameshift that knocked out the expression of OPA1. The sequences of the sgRNAs were: 5’-UAGAUUCAGCAGAUAAUUGA-3’ and 5’-GAAUUCAAAGCUCCUAAAGU-3’. MDA-MB-231 cells were electroporated (NEON electroporator, ThermoFisher) with the CRISPR RNP complex formed by mixing Cas9 protein (IDT) and sgRNAs (Synthego) *in vitro*. The cells were plated 48 hours after electroporation onto 96 well plates at a density of less than one cell per well and incubated for two weeks. Single cell clones were expanded and screened by genomic PCR using OPA1 primers. The loss of OPA1 expression in clones with homozygous deletion was further confirmed by western blot.

### *In vitro* irradiation and drug treatments

Cells were seeded and allowed to adhere and grow for 24-48 hours prior to treatments. Cells were irradiated using an XRAD320 Biological Irradiator (Precision X-Ray) with pre-calibrated doses. The dishes containing the cells were placed on a 1 cm thick PMMA-slap 50 cm from the source and irradiated with 320 kVp x-rays at 12.5 mA with 2.0 mm aluminum filter. MDA-MB-231 and SUM159pt cells were irradiated at 2, 4, 6, 8, 10, and 12 Gy (2.6 Gy/min). BCM-HCI-7821 PDXOs were irradiated at 8 Gy (2.6 Gy/min).

For MYLS22 treatments, MYLS22 powder (InvivoChem, V41464) was dissolved in DMSO (Sigma-Aldrich, D2650) by vortexing and sonication to make a 100 mM stock solution and stored at -80°C. Working concentrations (50 µM) were made in the appropriate complete growth medium for the cell line by vortexing and sonicating immediately after adding stock MYLS22 to media. Completely dissolved working concentrations were then added to cells.

### Measuring cell number and size

1 × 10^5^ cells were seeded in 6 cm dishes and left to adhere and grow for 36-48 hours prior to radiation. 48 hours following irradiation, cells were detached from the plate using trypsin (Corning 25-053-CI) and counted using ViaStain AOPI Staining Solution (Revity CS2-0106) and a Cellometer K2 (Nexcelom Bioscience). The number of viable cells and their mean diameters (d) in suspension were recorded. Mean diameters were then converted into mean volumes using the formula: V=(4/3)π(d/2)^3^.

### Longitudinal cell growth measurements

1 × 10^5^ cells were seeded in 6 cm dishes and left to adhere and grow. Cells were irradiated on the second day after seeding. Cells were then trypsinized and seeded at 1 × 10^3^ per well in a poly-D-lysine (Sigma-Aldrich, P6704) coated 96-well plate within 4 hours after irradiation. The plate was then loaded in the Incucyte S3 Live Cell Analysis instrument (Sartorius) and live-cell images were obtained every 12 hours using a 10x objective lens (5 images per well). Longitudinal cell growth was assessed by measuring cell culture well confluence analyzed using the Incucyte AI Confluence Analysis workflow in the Incucyte Base Software (Sartorius).

### Clonogenic assays

100-2000 cells were seeded in 6 well plates and allowed to adhere overnight. Each plate was irradiated at 0, 2, 4, 8, or 12 Gy then incubated for 8-9 days. The media was removed from the plates and cells were fixed and stained with a 1 to 4 mixture of crystal violet solution (Sigma-Aldrich, HT90132) and absolute ethanol (Fisher Scientific, BP2818500) for 15 minutes. Staining solution was removed from the wells, and the plates were washed by submerging the entire plate in fresh water 4 times. After washing, the plates were air dried overnight then stained colonies were counted. Plating efficiency (PE) was calculated for each dose as the number of colonies counted divided by the number of cells seeded. Surviving fraction (SF) was then calculated as PE at the given dose divided by PE of the unirradiated control from the same experiment. Survival data were fitted to the linear-quadratic model, SF = exp(−(αD + βD²)), where D is dose in Gy and α and β were constrained to be non-negative. Curves were compared using the extra sum-of-squares F test.

### Cell viability assay in cell lines

Cell viability following MYLS22 and irradiation treatments in MDA-MB-231 and SUM159pt cells was measured using CellTiter-Glo Luminescent Cell Viability Assay (Promega, G7572) according to the manufacture’s protocol. 1 × 10^3^ cells were seeded in 96 well plates and left to adhere and grow. MDA-MB-231 or SUM159pt cells were treated with 50 µM MYLS22 the day following seeding. The following day, MDA-MB-231 cells were irradiated at 4Gy and SUM159pt cells at 8 Gy. On the fifth day following irradiation, cells were removed from the incubator and allowed to come to room temperature for 15 minutes. CellTiter-Glo reagent was then added to each well in an equal volume to the current media in the well. Cells were then lysed by shaking at 700 rpm for 2 minutes at room temperature. Following lysis, 200 μl of each well was transferred to a white 96 well plate (Greiner Bio-One, 655073) and incubated for 10 minutes at room temperature protected from light. Luminescence was then measured using a BioTek Synergy LX plate reader.

### PDXO maintenance and cell viability measurements

Previously established^16,17^ patient-derived xenograft organoid (PDXO) model, BCM-HCI-7821, was thawed from a frozen stock and cultured in 200 μl domes of Matrigel matrix for organoid culture (Corning) in six-well plates (Genesee Scientific). PDXOs were covered with Advanced DMEM/F12 (Thermo Fisher) supplemented with 5% FBS, 10 mM HEPES (Thermo Fisher), 1X Glutamax (Thermo Fisher), 1 μg/ml hydrocortisone (Sigma-Aldrich), 50 μg/ml gentamicin (Genesee Scientific), 10 ng/ml hEGF (Sigma-Aldrich), and 10 μM Y-27632 (Selleck Chemicals) (TNBC PDXO medium). Prior to experimentation, PDXOs were authenticated through STR analysis, devoid of stromal cells, and were tested to ensure a stable doubling time.

4 × 10^5^ BCM-HCI-7821 PDXO cells were embedded in 200 µl domes of organoid Matrigel (Corning) in six-well plates (Genesee Scientific) and cultured in TNBC PDXO medium. Following thirteen days of growth, organoids grew to an average of 100 µm in diameter and were irradiated at a dose of 8 Gy. Seven days after IR, organoids were collected as previously described^18^ by dispase (50 units/mL) solution (*i.e.* dispase stock supplemented with 20% FBS and 10 mM Y-27632) treatment for 10-15 minutes at 37 °C. Organoids were then dissociated by incubating in TrypLE Express (Thermo Fisher, 12605010) supplemented with 10 µM Y-27632 and pipetting. Cells were washed in TNBC PDXO medium and counted using ViaStain AOPI Staining Solution (Revity CS2-0106) and a Cellometer K2 (Nexcelom Bioscience). The number of viable cells was recorded, and cells were pelleted and stored at -80°C.

### Proteomic profiling

2.5 × 10^5^ MDA-MB-231 cells were seeded in 10 cm plates and allowed to adhere and grow for 48hrs. Cells were then irradiated at 8 Gy. Cells were collected by washing cells on plates with cold PBS, scraping cells from the plate with cold PBS, pelleting the cells by centrifugation at 500 x g for 5 minutes, aspirating supernatant and snap freezing. Untreated cells were collected 3 days post seeding. Irradiated cells were collected two days and five days post-IR.

The cell pellets were lysed in 8M urea buffer, reduced/alkylated and digested using LysC and Trypsin proteases at 37°C overnight. The digested mixture was acidified with 1% final formic acid. The peptide desalting was done using the AssayMAP 5µl C18 cartridges on the Agilent AssayMAP Bravo system. Peptide concentration was determined by colorimetric assay (Pierce™ Quantitative Colorimetric Peptide Assay, Thermo Scientific) and samples were reconstituted at 100 ng/µL in 0.1% formic acid containing 0.015% DDM. LC-MS/MS analysis was performed on a nanoElute2 UHPLC system coupled online to a timsTOF Ultra 2 mass spectrometer (Bruker Daltonics) using a 25 cm × 75 µm × 1.5 µm PepSep column with a 60-min gradient.

Data were acquired in diaPASEF mode (m/z 400–1000; ion mobility 0.64–1.37 1/K0; 24 DIA windows; cycle time 0.96 s). Raw data were searched in library-free mode using DIA-NN (v2.2.0) against a human proteome FASTA (UniProtKB; 20,428 entries; accessed October 2024). Search parameters included trypsin/P digestion (up to 1 missed cleavage), variable oxidation (M) and N-terminal acetylation, fixed carbamidomethylation (C), and precursor FDR set at 1% with match-between-runs enabled. The DIA-NN precursor-level report file was processed through gpGrouper algorithm^19^ to obtain protein level quantification using the iBAQ approach. The Ms1.Normalised values were used to calculate iBAQ protein abundances, which were subsequently median normalized per sample then log2-transformed for statistical analysis. The data was further processed by removing proteins that were not detected or had a value of 0 in 50% or more of samples (1237/9171 proteins removed). Principal component analysis was run on the top 10% most variable proteins that were quantified in all samples.

Gene set variation analysis (GSVA) was conducted to derive sample-level pathway enrichment scores. Pathway gene sets were drawn from the KEGG_Legacy^20,21^ and MitoCarta3.0 MitoPathways^22^ collections. Enrichment scores were computed using the GSVA R/Bioconductor package^23^ with a Gaussian kernel and the maxDiff scoring scheme. Gene sets were filtered to those containing 5–500 measured genes and sets outside this range were excluded from scoring. One way ANOVA with Benjamini-Hochberg (BH) false discovery rate (FDR) correction applied was performed on GSVA scores from each collection the sorted by lowest FDR and the top 25 pathways were visualized in heatmaps. 135 out of 162 KEGG_Legacy pathways and 82 out of 102 MitoCarta3.0 pathways varied significantly at FDR <0.05.

To determine changes in individual protein expression between treatment groups, normalized log2-transformed iBAQ protein abundances were used. Day 2 vs. untreated and day 5 vs. untreated pairwise group comparisons were calculated using moderated t-tests implemented in limma^24^ with BH FDR correction applied. The signed-log10(FDR) values for oxphos proteins in day 5 vs. untreated (x-axis) and day 2 vs. untreated (y-axis) were plotted as a two-way volcano plot. ETC proteins complex I-V are denoted in different colored dots. Mitochondrially encoded proteins are denoted with a red ring.

### Metabolomic profiling

2.5 × 10^5^ MDA-MB-231 cells were seeded in 10 cm plates and allowed to adhere and grow for 48hrs. Cells were then irradiated at 8 Gy. Cells were collected by washing cells on plates with PBS, trypsinization (Corning, 25-053-CI), trypsin neutralization and collection with growth medium, and pelleting the cells by centrifugation at 170 x g for five minutes. Cell pellets were then washed in PBS and counted 3 times for accuracy. 300K cells were transferred to microcentrifuge tubes and pelleted at 500 x g for five minutes then snap frozen. Untreated cells were collected three days post seeding. Irradiated cells were collected two days and five days post-IR.

Central carbon metabolites, including amino acids, were extracted from pellets using a previously described liquid–liquid extraction method^25–28^. A pooled QC sample was included throughout the mass spectrometry acquisition to monitor instrument performance and data reproducibility. Chromatographic separation was performed using hydrophilic interaction liquid chromatography (HILIC) on an XBridge Amide column (3.5 μm, 4.6 × 100 mm; Waters) operated in positive electrospray ionization (ESI+) mode. The mobile phases consisted of water containing 0.1% formic acid (Mobile Phase A) and acetonitrile containing 0.1% formic acid (Mobile Phase B)^29^.

Redox metabolites were extracted from cultured cells by adding 100 μL of ice-cold extraction solvent consisting of methanol: acetonitrile: water (4:4:2, v/v/v) containing 0.1% formic acid. Cell pellets were disrupted by probe sonication, followed by centrifugation at 15,000 rpm for 10 min at 4°C. Subsequently, 90 μL of the supernatant was transferred to a clean microcentrifuge tube and dried under vacuum using the aqueous mode for approximately 40 min. The dried extracts were reconstituted in 100 μL of methanol: water (1:1, v/v), vortexed thoroughly, and 10 μL was injected for LC–MS analysis. Chromatographic separation was performed on a Phenomenex Luna® NH₂ column (150 × 2.0 mm, 3 μm particle size, 100 Å; Phenomenex, Torrance, CA, USA) using a binary UHPLC system. The mobile phases consisted of 100% acetonitrile (Mobile Phase A) and 5 mM ammonium acetate in water (pH 9.0; Mobile Phase B). The gradient program was as follows: 10% B from 0–20 min, increased to 90% B from 20–25 min, returned to 10% B at 26 min, and maintained at 10% B until 34 min for column re-equilibration. The flow rate was maintained at 0.25 mL/min throughout the analysis.

Glycolysis, pentose phosphate pathway, and TCA metabolites (carbohydrate panel) were extracted from cultured cells as previously described^26,28,30–32^. A pooled QC sample was prepared and analyzed throughout the analytical run as a quality control (QC) sample. Chromatographic separation of TCA cycle and glycolytic intermediates was performed using a Luna NH₂ column (3 μm, 100 Å; Phenomenex). The mobile phases consisted of 5 mM ammonium acetate in water (pH 9.9) (Mobile Phase A) and acetonitrile (Mobile Phase B).

Targeted metabolite analysis was performed using an Agilent 1290 Infinity UHPLC system coupled to an Agilent 6495B Triple Quadrupole Mass Spectrometer (Agilent Technologies, Santa Clara, CA). Metabolites were detected using the Multiple Reaction Monitoring (MRM) acquisition mode. TCA cycle metabolites were analyzed in negative electrospray ionization (ESI−) mode, whereas central carbon and redox metabolites were analyzed in positive electrospray ionization (ESI+) mode^31,32^. Raw data were processed using Agilent MassHunter Quantitative Analysis software. Peak integration was manually inspected and curated, when necessary, on a sample-by-sample basis to ensure accurate quantification.

Metabolite peak areas were normalized to the corresponding internal standards and log2-transformed for statistical analyses. Principal component analysis (PCA) was done using all metabolites for each panel. Normalized log2 abundance values were scaled for each metabolite and plotted as a heatmap.

Differential metabolites between experimental groups were identified using moderated t-tests using limma^24^, followed by Benjamini–Hochberg correction for multiple hypothesis testing. Log2 fold changes between groups were used to determine the magnitude of change.

### Animal Studies

This study was carried out in accordance with the *Guide for the Care and Use of Laboratory Animals* from the National Institutes of Health (NIH) IACUC. The protocol was approved by the IACUC at BCM (protocol AN-8243). Mice were ethically euthanized when they reached defined study end points. Euthanasia was conducted as recommended by the Association for Assessment and Accreditation of Laboratory Animal Care International.

MDA-MB-231 cells in culture were detached using trypsin, washed in ‘231 growth medium’ (see Cell culture and authentication section) and counted by AOPI staining with a Cellometer K2 (Nexcelom Bioscience). Viable cells were resuspended in a 1:1 mixture of ‘231 growth medium’ and Matrigel (Corning, 354234) at a concentration of 5 × 10^7^ cells/mL. 20µl of tumor cell suspension was then injected unilaterally into the fourth mammary fat pad of five-week-old SCID/bg mice (C.B-17/IcrHsd-*Prkdc^scid^Lyst^bg-J^*, Inotiv).

Tumor length and widths were measured using digital calipers and tumor volumes were calculated: V=(1/2)LW^2^. After tumors reached an average volume of approximately 350 mm^3^, mice were irradiated. All mouse irradiations were performed using the x-ray source irradiator (XRAD320), a self-contained x-ray system (Precision X-Ray). The system is fitted with an adjustable/variable collimator [0 to 20cm x 20cm (at 50 cm SSD) x-ray field size]; tube specs: Max; potential: 320kVp that delivers 3 Gy/min at 12.5mA, 50cm SSD (HVL≈ 1mm Cu). The XRAD320 is equipped with an internal dose measurement and control system. The XRAD320 is calibrated yearly by the company to confirm that internal dosimeter matches external ion chamber measurements. Animals were anesthetized before and during irradiation using an isoflurane/oxygen mixture. The right 4th mammary gland was outlined, and the variable collimator was adjusted to a field of 1.5 X1.5cm to target only the tumor while the rest of the body was covered with 1cm thick lead sheet, to ensure shielding of the body from scatter radiation. A single dose of 10Gy was delivered to the targeted radiation field. Seven days following IR, three mice from each group were euthanized and their tumors were removed. The remaining mice were followed until an ethical endpoint was reached (day 12 for naïve mice and day 19 for IR mice). Tumor pieces were snap frozen in liquid nitrogen and stored at -80°C as well as processed into formalin-fixed paraffin-embedded (FFPE) blocks.

### Immunohistochemistry (IHC) staining and analysis

Tumor pieces were fixed in 10% neutral buffered formalin for 48 hours, washed three times in PBS, and stored at 4°C in 70% ethanol. Fixed tumors were then embedded in paraffin creating FFPE blocks. FFPE blocks were cut into 3 µm sections and placed on slides. Routine hematoxylin and eosin (H&E; Epredia, 72711 and 71311) staining was conducted. IHC was conducted with antibodies against Ki67, CC3, phospho-histone H3 (pHH3), p21, and SOD2 (**Supplementary Table 1**). Antigen retrieval was done with 0.01M citrate buffer, pH6 (for pHH3) or 0.1M Tris-HCL, pH 9 (for all other proteins) for 15 min at full pressure (above 90 °C) with a pressure cooker. Stained slides were imaged on a Nikon Eclipse Ci microscope with a 40X objective. Images were further processed using ImageJ.

### Transmission electron microscopy and analysis

Cell cultures were fixed, processed and embedded in situ to prevent morphological damage, as follows. Cells were fixed in Karnovsky’s fixative, post-fixed and stained in a solution of 1% osmium tetroxide and 1.5% potassium ferricyanide for one hour, stained in 3% uranyl acetate overnight at 4°C, dehydrated in a series of ethanol solutions of increasing concentrations, and embedded in Spurr’s resin. Ultra-thin sections of all samples were cut using a Leica EM UC7 ultramicrotome and imaged using a JEOL JEM-1400Flash TEM equipped with an AMT NanoSprint15 sCMOS camera. More than 55 images per sample were taken at direct magnifications from x800–5000. Mitochondria from a minimum of 30 cells per sample were annotated and cristae were scored from 0-4 as previously described^33^ using ImageJ software. Briefly, a score of 4 indicated >75% of a mitochondrion contained regular cristae, 3 indicated >75% of a mitochondrion contained irregular cristae, 2 indicated 50-75% of a mitochondrion contained cristae, 1 indicated <50% of a mitochondrion contains cristae, and 0 indicated a mitochondrion was devoid of defined cristae.

### Fluorescence microscopy imaging and analysis

The mitochondria-specific fluorescent dye MitoTracker CMXRos (Thermo Fisher Scientific, M7514) was used to monitor mitochondrial morphology as described previously^11^. Briefly, cells were seeded on coverslips (Thorlabs, CG15XH1) coated with Poly-D-lysine (Sigma-Aldrich, P6407) and irradiated at 8 Gy (MDA-MB-231) or 12 Gy (SUM159pt) on the second day following seeding. Coverslips were stained, fixed and mounted 2, 5, 7, 12, or 15 days after irradiation. On these days, cells were treated with fresh growth media containing 100 ng/mL MitoTracker CMXRos (Invitrogen, M7512) at 37°C for 15–30 minutes. Cells then were washed three times with growth media and fixed with pre-warmed media containing 3.7% formaldehyde solution for 15 minutes at 37°C. After fixation, cells were washed three times with PBS, permeabilized with 0.2% Triton X-100 for 10 minutes at room temperature, then stained with 300 nM DAPI in PBS (Invitrogen, D3571). Coverslips were mounted on glass slides with ProLong mounting Medium (Invitrogen, P36982).

Fluorescence imaging was carried out on a Zeiss LSM880 with Airyscan confocal microscope using a 63X oil objective. MitoTracker was excited using the 561 nm laser and emitted light around 599 nm was captured. DAPI was excited using the 405 nm laser and emitted light around 461 nm was captured. Z-stacks of 4 to 10 slices were generated for each image. Raw images were processed and z-stacks compressed into maximum projections using ImageJ. 15-40 cells per sample were imaged and analyzed. Mitochondria shape and number were analyzed by the MAT-LAB based macro, MicroP according to the author’s instructions^34^. Mitochondria identified as “small globe” or “large globe” were classified as “fragmented”. Mitochondria identified as “simple tube”, “twisting tube”, “donut”, or “branching tube” were classified as “elongated”. Area per cell was calculated by summing the area of each mitochondrion in an image and dividing the total by the number of cells in the image.

### DNA extraction and mtDNA quantification

Total DNA was extracted from snap frozen tumor chunks and cell pellets using the Mag-Bind® Blood & Tissue DNA HDQ 96 Kit (Omega Bio-Tek, M6399-00) according to manufacturer’s instructions. Purified DNA samples were quantified by NanoDrop 2000 (Thermo Scientific). Quantitative PCR was performed as previously described^11^ with the Universal SYBR Green Supermix (Bio-Rad, 1725121), 2 ng of DNA, and 1 µM of primers (Sigma-Aldrich) recognizing ND1, ND6, RGPD1, and FUNDC2P2 **(Supplemental Table 2)** with cycling conditions: 95 °C for 10 minutes followed by 40 Cycles of: 95 °C for 15 seconds 64 °C for 1 minute. Relative mtDNA copy number was calculated using the relative 2^−ΔΔCt^ method where averaged Ct values of the mitochondrial genes (ND1 and ND6) were compared to the averaged Ct values of the nuclear genes (RGPD1 and FUNDC2P2).

### Western blotting

Whole cell lysates were generated by lysing cells with RIPA buffer (Thermo Scientific, 89901) supplemented with 1 tablet of protease inhibitor cocktail (Roche, 11836153001) and 1 tablet of PhosSTOP™ (Roche, 4906837001) per 10 mL of RIPA. After lysis with supplemented RIPA buffer, cell debris was removed by centrifugation at 20,000xg for 20 minutes at 4°C and soluble proteins were collected. Protein samples were quantified by Pierce BCA assay (Thermo Scientific, 23222). Protein samples were diluted in 4X sample buffer (Bio-Rad, 1610747) with 10% beta-mercaptoethanol (Sigma-Aldrich, M6250) and denatured by heating at 100°C for 10 minutes. Diluted protein samples were separated on pre-cast 4-20% gradient (Bio-Rad, 5671095) or 7.5% SDS-polyacrylamide gels (Bio-Rad, 5671024) then transferred to a nitrocellulose membrane (Bio-Rad, 1704271) using the Trans-Blot Turbo system (Bio-Rad). Membranes were then stained with Ponceau S (Tocris, 5225) for 10 minutes at room temperature then destained with DI water and imaged to assess transfer quality. After blocking with EveryBlot Blocking Buffer (Bio-Rad, 12010020) for 10 minutes at room temperature, membranes were incubated with primary antibodies **(Supplemental Table 1)** diluted in Blocking Buffer overnight at 4°C on a rocker, washed with 0.1% Tween 20 (Sigma-Aldrich, P9416) in PBS (PBST), then incubated with HRP-conjugated secondary antibody diluted in Blocking Buffer for 1 hour at room temperature. Blots were washed with PBST and then detected using Clarity Western ECL Substrate (Bio-Rad, 1705060) or SuperFemto ECL Chemiluminescence kit (Vazyme, E423-02). Images were acquired on a ChemiDoc Touch Imaging System (Bio-Rad) and analyzed using ImageLab software v.6.1 (Bio-Rad).

### Metabolic flux analysis

Mito Stress Tests were performed using a Seahorse XFe96 extracellular flux analyzer (Agilent) to assess mitochondrial function and other metabolic changes after irradiation. 1 × 10^5^ cells were seeded in 6cm dishes then irradiated at 8 Gy (MDA-MB-231) or 12 Gy (SUM159pt) on the second day following seeding. At the time of the assay, cells were reseeded at 50,000 cells per well in Poly-D-lysine (Sigma-Aldrich, P6407) coated XFe96 cell culture microplates (Agilent, 103792-100) in Seahorse XF RPMI medium pH 7.4 (Agilent, 103576-100) supplemented with 10 mM glucose (Agilent, 103577-100), 1 mM sodium pyruvate (Agilent, 103578-100), and 2 mM L-glutamine (Agilent, 103579-100). A minimum of four technical replicates were run for each biological condition. The plates were centrifuged for 1 minute at 1000 rpm, then incubated for 45 minutes to 1 hour at 37°C in a CO_2_-free incubator. During this time, drugs from the Mito Stress Test Kit (Agilent, 103015-100) including oligomycin (1.5 μM), FCCP (1 μM), and rotenone/antimycin A (0.5 μM), were loaded into the injection ports of the XFe96 sensor cartridge (Agilent, 103792-100), which was then loaded into XFe96 analyzer for calibrating. After calibration, the XFe96 microplate was loaded into the Seahorse bioanalyzer for analysis. Upon completion of the assay, OCR values were assessed for outliers using the Grubb’s outlier test and wells were eliminated from analysis. Basal respiration, maximal respiration, mitoATP production, glycoATP production, non-mitochondrial OCR, proton leak, spare respiratory capacity, coupling efficiency, and basal ECAR for each run were all calculated using seahorseanalytics.agilent.com. Technical replicate values for each run were averaged to generate each biological replicate value.

### ROS measurement

2.5 × 10^5^ cells were seeded in 10cm plates then irradiated at 4 and 8 Gy (MDA-MB-231) or 8 and 12 Gy (SUM159pt) on the second day following seeding. 24hr after irradiation, cells were collected by trypsinization and resuspended in “Assay Buffer” consisting of Seahorse XF RPMI medium pH 7.4 (Agilent, 103576-100) supplemented with 5 mM glucose (Thermo Fisher, A2494001), 2000 mg/L sodium bicarbonate (Sigma-Aldrich, S8761), 2 mM L-glutamine (Sigma-Aldrich, G7513), and 10% fetal bovine serum (R&D, S11550). Cells were counted then diluted to 0.4-1 × 10^6^ cells per mL and aliquoted into Falcon round bottom polystyrene tubes. 2′,7′-dichlorodihydrofluorescein diacetate (DCFDA) powder (Lumiprobe, 2247) was dissolved in DMSO (Sigma-Aldrich, D2650) by vortexing to make 5 mM stock solution, aliquoted, and stored at -20°C. 40 mM working solution was made by diluting stock DCFDA in Supplemented Buffer. 40 mM working solution or DMSO control solution was then added to cells in Falcon tubes in a 1 to 1 ratio for a final staining concentration of 20 mM. The stained cells were then incubated at 37°C with 5% CO_2_ for 1-1.5 hours. Dead cell marker Sytox Blue (Thermo Fisher, S34857) was then added to each sample and cells were analyzed with a BD FACSCanto II flow cytometer using the V450 channel to detect Sytox Blue and B530 channel to detect DCFDA signal. Data was collected using the BD Divas software and analyzed using FlowJo 10 software. Single channel, positive, and negative controls were used for each experiment. Data was gated for single and live cells then geometric mean fluorescence intensity of DCFDA signal was calculated for each sample.

### Analysis of mitochondrial pathway activation in murine p53 null tumor microarray data

Oh *et al.*^35^ published microarray data from the T11, 2225L, and 2250L murine breast cancer models 4, 8, 12, 24, and 48 hours after irradiation in vitro. Each timepoint had a matched non-irradiated control. For each timepoint, a fold change value for every gene in the microarray was calculated by dividing the gene’s level in the irradiated sample by its level in the non-irradiated control. ssGSEA^36^ was conducted on these fold change values to determine the NES and associated p value for MitoCarta 3.0 pathways^22^ at each timepoint. NES values were then represented as a heatmap for each timepoint for each pathway, with special indication of those NES scores whose corresponding p value was less than 0.05 and thus met significance criteria.

### Correlation of RNA expression with irradiation resistance in breast cancer cell lines

Yard *et al.*^37^ published radiation resistance scores (AUC) for 533 cell lines, 28 of which were breast cancer (BRCA) cell lines. Baseline RNA-seq expression for each of these 28 BRCA cells lines were obtained from the DepMap resource (www.DepMap.org) as log2(TPM+1) values for protein coding genes. Spearman correlations between expression and radiation AUC were computed per gene. Genes were then nranked by signed -log10(p), with the sign taken from the Spearman coefficient (rho > 0, positive; rho < 0, negative). The ranked gene list was then analyzed by GSEA^21^ was performed against KEGG pathways^20^ using WebGestaltR v0.4.6^38^ with default settings. Pathways with FDR < 0.05 are plotted by normalized enrichment scores (NES).

### Statistical analysis

Unless otherwise noted in figure legends, the data are shown as mean ± standard error (SEM), and all experiments were repeated at least three times. Statistical tests were conducted using GraphPad Prism or R software. Outlier samples were identified by Grubb’s test in Prism and removed. All data meet normal distribution and have uniform standard deviations, unless otherwise noted.

## RESULTS

### TNBC cells surviving radiation treatment enter a plastic residual state that largely reverts upon regrowth

To model residual disease persisting after ionizing radiation (IR) treatment, we treated human TNBC cells with a single dose of IR, leaving behind a population of residual cells that would invariably regrow after initial ‘regression’ of cells. The effective IC_70_ doses for MDA-MB-231 and SUM159pt cells were 8 Gy and 12 Gy, respectively, two days after treatment (**Figure S1A-B**). Accordingly, clonogenicity of each cell line decreased in a dose-dependent manner, with MDA-MB-231 cells being more sensitive than SUM159pt cells (**Figure S1C**).

We noted striking changes in cellular morphology in MDA-MB-231 cells, with many cells adopting giant cell-like features (**Figure 1A-C**). Giant cell-like morphology arose as early as two days after treatment with 8 Gy and became more exaggerated until the population nadir was reached (*i.e.,* the minimum number of cells persisting following IR) five days after treatment. Spontaneous cell regrowth began 7-10 days after IR and continued for 5-7 more days until the plate became confluent. Notably, cells largely reverted to their pre-treatment morphology as the population regrew from the residual state (**Figure 1A**). IC_70_ treatment of SUM159pt cells did not yield notable changes in cell morphology but did elicit a trend towards larger cell volumes with a similar timeline of cell regression and regrowth as observed in MDA-MB-231 cells (**Figure S1D-F**).

**Figure 1.**
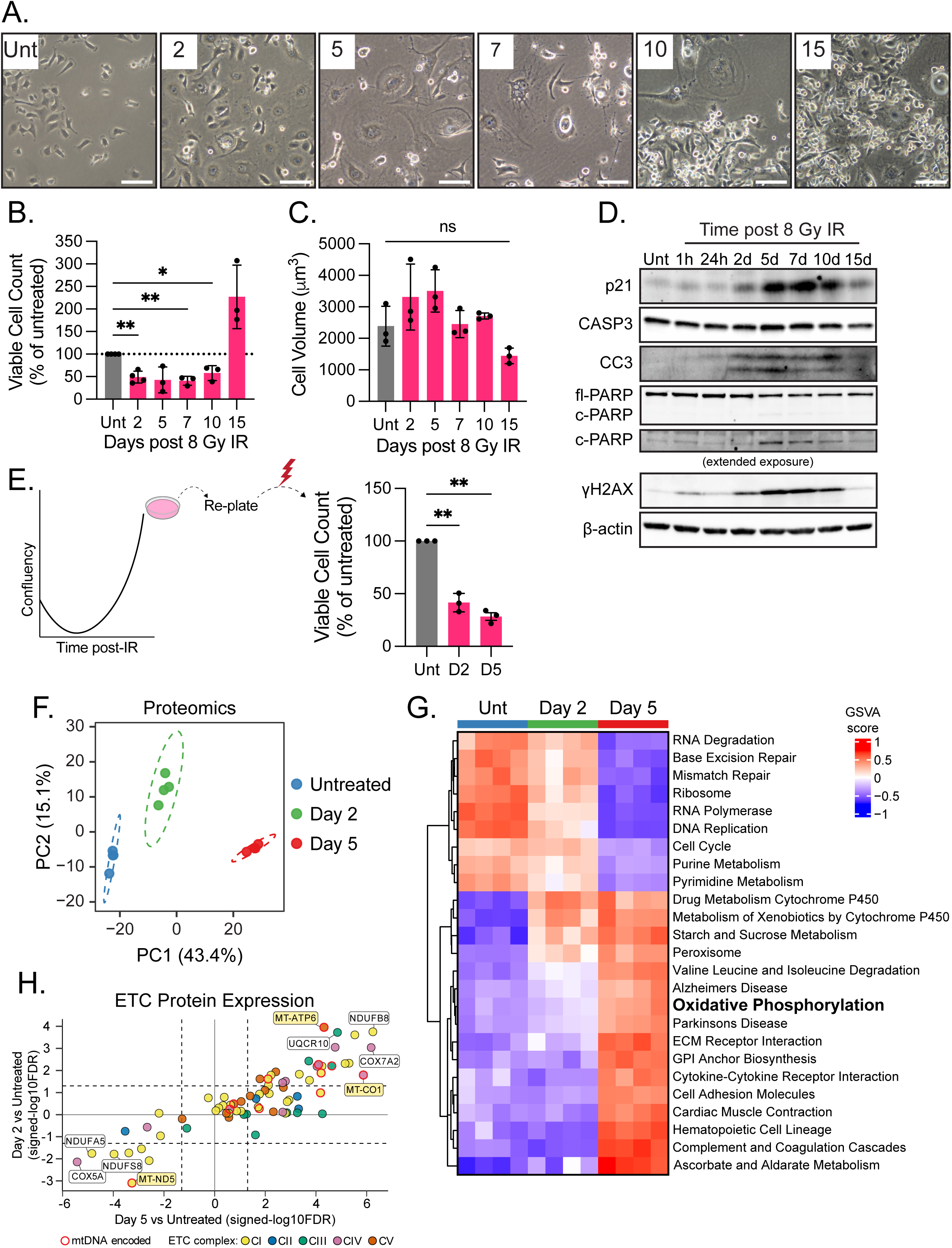
TNBC cells surviving radiation treatment enter a plastic residual state that largely reverts upon regrowth. (A) Brightfield images of MDA-MB-231 cells taken longitudinally after 8 Gy of IR. Scale bar is 100 µm. (B) Relative viable MDA-MB-231 cells counted after AO/PI staining 2, 5, 7, 10, and 15 days after 8 Gy IR. Counts were normalized to untreated cells collected at day 2 post-IR. **P<0.01, *P<0.05 by one sample t-test compared to 100, n≥3. (C) Mean cell volumes calculated from mean diameters of viable cells 2, 5, 7, 10, and 15 days after 8 Gy IR. No significant change by ANOVA. (D) Representative immunoblots of quiescence, apoptosis, and DNA damage-related proteins in MDA-MB-231 whole cell lysates collected at days 2, 5, 7,10, and 15 post 8 Gy IR. (E) Relative viable regrown MDA-MB-231 cells counted by AO/PI staining 2 and 5 days after 8 Gy IR. Counts were normalized to untreated cells collected at day 2 post-IR. **P<0.01 by one sample t-test compared to 100, n=3. (F) Principal component analysis (PCA) of mass spectrometry proteomics of MDA-MB-231 cells at days 2 and 5 post 8 Gy IR. (G) Heatmap displaying KEGG pathways by gene set variation analysis (GSVA). Pathways were sorted by ANOVA and the top 25 most variable were displayed, n=4. (H) Two-way volcano plot displaying signed-log10(FDR) values of ETC proteins comparing day 5 vs. untreated (x-axis) and day 2 vs. untreated (y-axis). Significance cutoff lines are at |1.3|.

Giant cell morphology is often associated with stress-induced cellular quiescence^39^. Concomitant with the noted cellular regression and regrowth, immunoblotting revealed similar temporal dynamics of p21 and apoptosis markers cleaved caspase 3 (cCASP3) and cleaved Poly (ADP-ribose) polymerase (cPARP) (**Figure 1D; S1G**). DNA double-strand breaks (DSB) were elevated, as manifested by levels of S139 phosphorylated-histone H2A.X (γH2AX) as early as one-hour post-IR, which were resolved by 24hr, in alignment with the well-established temporal dynamics of DNA DSB repair following IR^40^. DSBs again accumulated starting on day 2 and diminished as cells regrew, suggesting indirect effects of IR led to gradual accumulation of DSBs after their initial resolution (**Figure 1D; S1G**). Further, we observed significant, dose-dependent elevation of ROS, an additional hallmark of radiation exposure, in both cell lines 24 hours after IR treatment (**Figure S1H-I**). Functionally, both cell lines retained radio-sensitivity once cells re-grew (**Figures 1E, S1J)**, affirming that survival of the radio-residual population reflects a transient adaptive state rather than a fixed, heritable feature.

Mass spectrometry proteomics profiling of MDA-MB-231 cells treated with 8 Gy of IR revealed a progressive rewiring of the proteome (**Figure 1F**). Gene Set Variation Analysis (GSVA) supported the finding that ‘radio-residual’ cells slow proliferation, perhaps adopting a quiescent phenotype as DNA replication, DNA repair, RNA metabolism, and ribosome pathways were significantly repressed following IR (**Figure 1G; Supplementary Table 3**). In contrast, IR induced many metabolic pathways including oxphos, amino acid metabolism, and fatty acid metabolism (**Figure 1G**). To gain deeper insights into metabolic rewiring induced by IR, we conducted GSVA analysis of MitoCarta 3.0 pathways^22^ (**Figure S2; Supplementary Table 4)**. Significantly altered MitoCarta pathways in post-IR cells fell under four main categories; metabolism, oxphos subunits, mitochondrial dynamics and surveillance, and mitochondrial central dogma (**Figure S2**). We noted most electron transport chain (ETC) proteins had higher expression in day 5 versus naïve cells, including those encoded in mtDNA as well as nDNA and encompassing all five complexes (**Figure 1H; Supplementary Table 5**). Together, these results provide evidence that following IR, TNBC cells can enter a ‘radio-residual’ state characterized by dynamic DNA damage and stress responses, metabolic rewiring, and quiescence.

### Residual cells surviving irradiation have elevated mitochondrial content and function

In light of the extensive upregulation of ETC protein levels observed by proteomics, we performed longitudinal MitoStress Tests to measure mitochondrial function. Basal and maximal oxygen consumption rates (OCR, *i.e.* oxphos) were significantly increased as early as 24 hours and continued to rise until five days following IR, the most ‘regressed’ residual time point (**Figure 2A-D; S3A-B**). As cells regrew, basal OCR levels dropped significantly in MDA-MB-231 cells, reverting to pre-treatment levels as plates became confluent (**Figure 2A**). Basal OCR tended to decrease in SUM159pt cells as they regrew but did not fully revert to pre-treatment levels (**Figure 2B**). MDA-MB-231 cells exhibited significant elevation, followed by reversion of maximal OCR and spare respiratory capacity (SRC) following IR (**Figure 2C; S3C**). In contrast, SUM159pt cells exhibited progressive elevation of maximal OCR and SRC throughout regression as well as regrowth (**Figure 2D; S3D**). Mitochondrial ATP production followed a similar temporal pattern to basal OCR in both cell lines (**Figure 2E-F**). Extracellular acidification rate (ECAR, *i.e.,* glycolysis) and coupling efficiency were largely unchanged following IR (**Figure S3E-H**). Non-mitochondrial OCR and proton leak were also transiently elevated in residual cells in both cell lines (**Figure S3I-L**), suggesting elevation of oxidative stress. Levels of the mitochondrial antioxidant superoxide dismutase 2 (SOD2) were increased at day 5 following IR and were sustained through regrowth in both cell lines (**Figure S4-B**). Together, these findings indicate that cells surviving IR specifically upregulate mitochondrial ATP production, oxphos rate, and oxphos capacity most strongly in the residual state, a phenotype that largely reverted as cells regrew.

**Figure 2.**
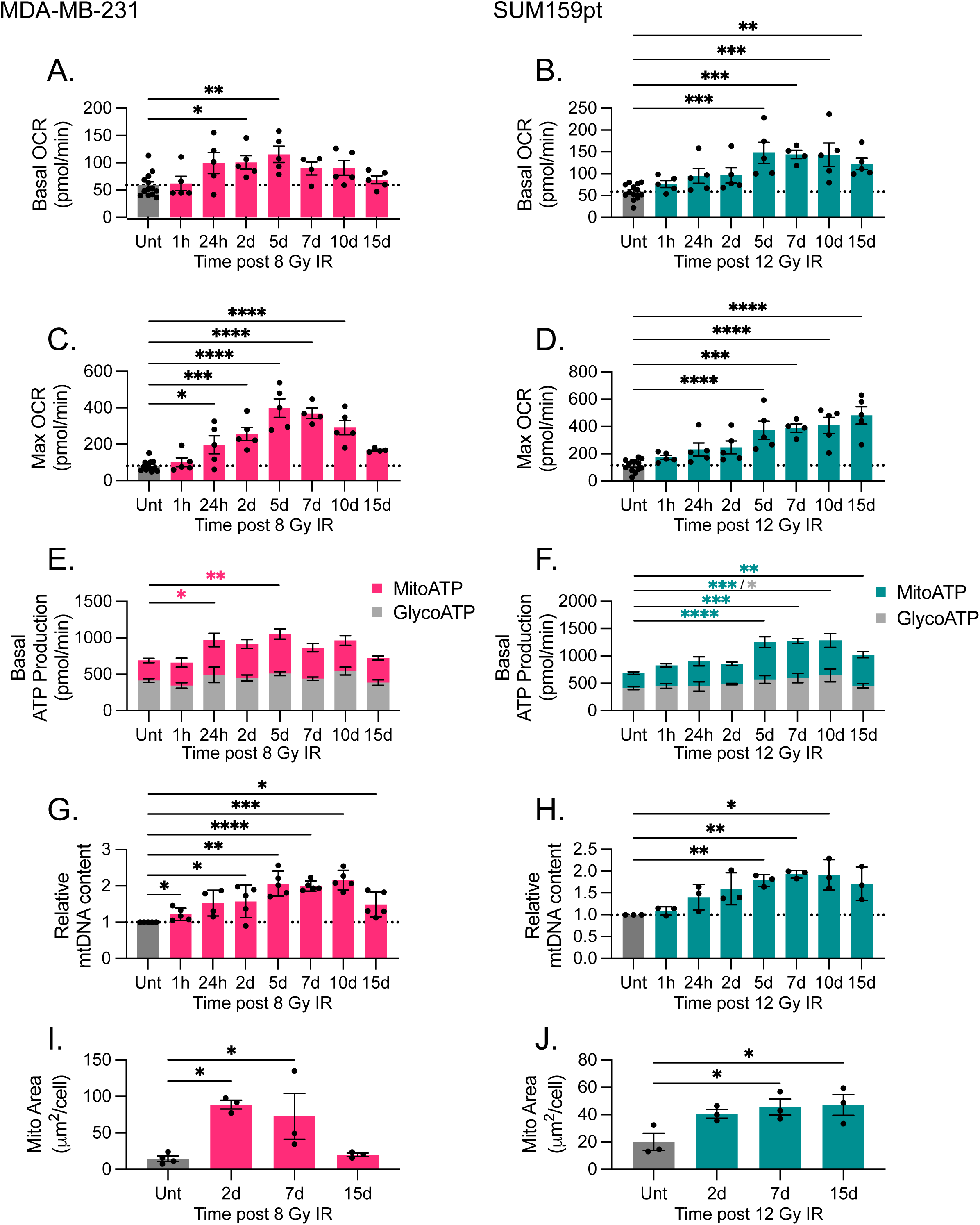
Residual cells surviving radiation have elevated mitochondrial content and function. (A) Quantified basal oxygen consumption rate (OCR) from Seahorse Mito Stress tests of MDA-MB-231 and (B) SUM159pt cells 1 hour, 24 hours, and 2, 5, 7, 10, and 15 days post-IR. (C) Maximal OCR in MDA-MB-231 and (D) SUM159pt cells. (E) Basal ATP production from mitochondria (MitoATP) and glycolysis (GlycoATP) in MDA-MB-231 and (F) SUM159pt cells. ****P<0.0001, ***P<0.001, **P<0.01, *P<0.05 by Dunnet’s multiple comparisons test following one-way ANOVA, n≥4. (G) mtDNA content of 8 Gy IR MDA-MB-231 and (J) 12 Gy IR SUM159pt cells normalized to untreated cells in each experiment. ****P<0.0001, ***P<0.001, **P<0.01, *P<0.05 by one sample t-test compared to 1, n≥3. (I) Quantification of mitochondrial area (MitoTracker staining) per cell by in MDA-MB-231 cells and (J) SUM159pt cells. Data are presented as biological replicates (n≥3) of at least 15 cells per experiment. *P<0.05 by Dunnet’s multiple comparisons test following one-way ANOVA

We observed significant elevation of mtDNA levels matching the temporal trends of oxphos changes (**Figure 2G-H**). Concordantly, using MitoTracker Red staining, we observed a significant increase in mitochondrial area per cell in radio-residual cells which returned to baseline in regrown MDA-MB-231 cells but remained elevated in SUM159pt cells (**Figure 2I-J**). However, these changes were not associated with elevation of mitochondrial biogenesis signatures in the proteomics profiling data (**Supplementary Table 5**), suggesting mitochondrial mass may be regulated by additional mechanisms following IR.

We broadened these findings by treating an orthotopic TNBC patient-derived xenograft (PDX)-derived organoid (PDXO) model, BCM-HCI-7821^16,17^, with IR. Treatment with 8 Gy led to increased mtDNA content, nuclear DNA DSBs, and p21 levels, consistent with our findings in human cell lines (**Figure 3A-C**). We extended our studies *in vivo* using an MDA-MB-231 orthotopic xenograft. Mammary tumors were focally treated with a single 10 Gy dose of IR, resulting in sustained tumor stasis (**Figure 3D**). Congruent with our *in vitro* findings, irradiated tumors had elevated mtDNA copy number (**Figure 3E**), with tumor cells exhibiting larger size and aberrant morphologies seven days after IR (**Figure 3F**). While we observed no changes in Ki67 or CC3 staining, we did find fewer phosphorylated histone H3 (pHH3)-positive cells and more p21 positive cells in irradiated tumors compared to naïve tumors, consistent with the observed tumor stasis (**Figure 3D&F**). These studies provide *in vitro* and *in vivo* evidence of lR-induced tumor cell and mitochondrial adaptations in surviving TNBC cells.

**Figure 3.**
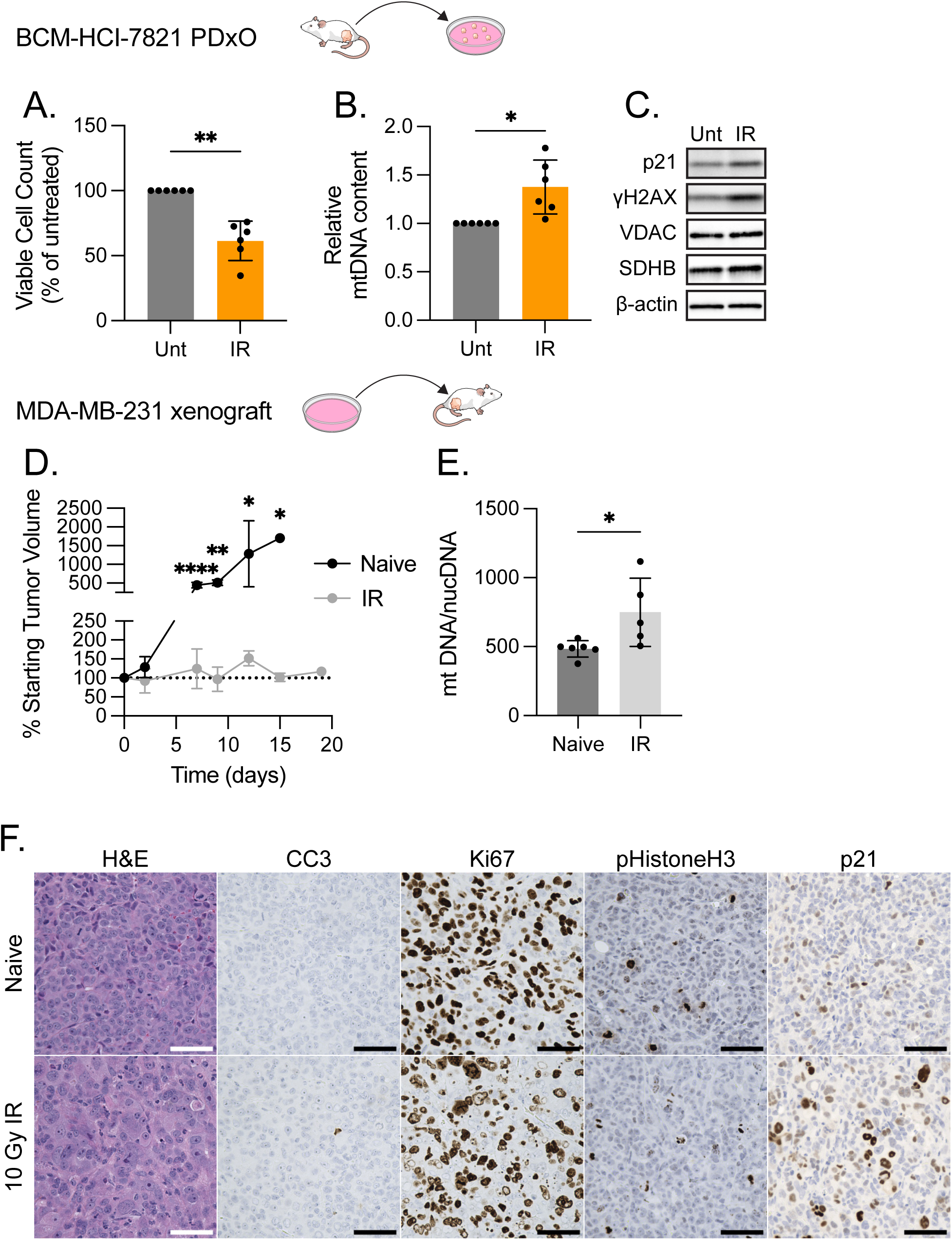
3D and *in vivo* models have elevated mitochondrial content following IR. (A) Relative viable cells counted after AO/PI staining and (B) mtDNA content of dissociated BCM-HCI-7821 PDXOs 7 days post 8 Gy IR. Counts were normalized to untreated cells in each experiment. **P<0.01, *P<0.05 by one sample t-test compared to 100 and 1, respectively. n=6 (C) Representative immunoblots in untreated and 8 Gy IR BCM-HCI-7821 PDXO cell lysates. (D) Percent change in MDA-MB-231 orthotopic xenograft tumor volumes following 10 Gy of focal IR. (E) mtDNA content of naïve and IR tumors. *P<0.05 by unpaired t-test, n≥5. (F) Representative H&E and IHC images of naïve and IR tumors collected at day 7 post-IR. Scale bar is 50 µm.

### Heightened TCA cycle and antioxidant responses accompany oxphos elevation in radio-residual cells

To further explore metabolic rewiring following IR of TNBC cells, we performed three targeted mass spectrometry metabolomic panels on the same irradiated MDA-MB-231 samples collected at day 2 and day 5 post-IR that were proteomically profiled. In the “central carbon” and “redox” panels, day 2 and day 5 samples clustered together while in the “carbohydrate metabolism” panel, day 2 and untreated samples clustered closely (**Figure S5A-C**), suggesting that alteration of central carbon and redox metabolism may precede rewiring of carbohydrate metabolism. We observed increases in many free amino acids and nucleotide metabolites following IR (**Figure S5D-F; Supplementary Table 6**), consistent with the observed reductions in “ribosome”, “RNA Degradation”, “RNA Polymerase”, “DNA Replication”, “purine metabolism”, and “pyrimidine metabolism” proteomic pathways (**Figure 1G**), perhaps attributable to reduced cell growth and proliferation.

To gain a better understanding of how metabolic rewiring may be related to increased oxphos, we integrated proteomics and metabolomics data focusing on carbohydrate metabolism. We observed an increase in most glycolytic intermediates and their cognate enzymes at day 5 post-IR (**Figure 4A**). Importantly, lactate was not significantly increased, suggesting the increase in glycolysis served to fuel the TCA cycle rather than lactic acid fermentation, in agreement with our prior findings in chemotherapy-treated TNBC cells^11^. Concordantly, we observed an increase in TCA cycle enzyme protein levels and a reduction in reactants from NAD(P)H producing reactions (*i.e.*, citrate/isocitrate and malate) suggesting increased TCA cycling at five days post-IR (**Figure 4A-B**).

**Figure 4.**
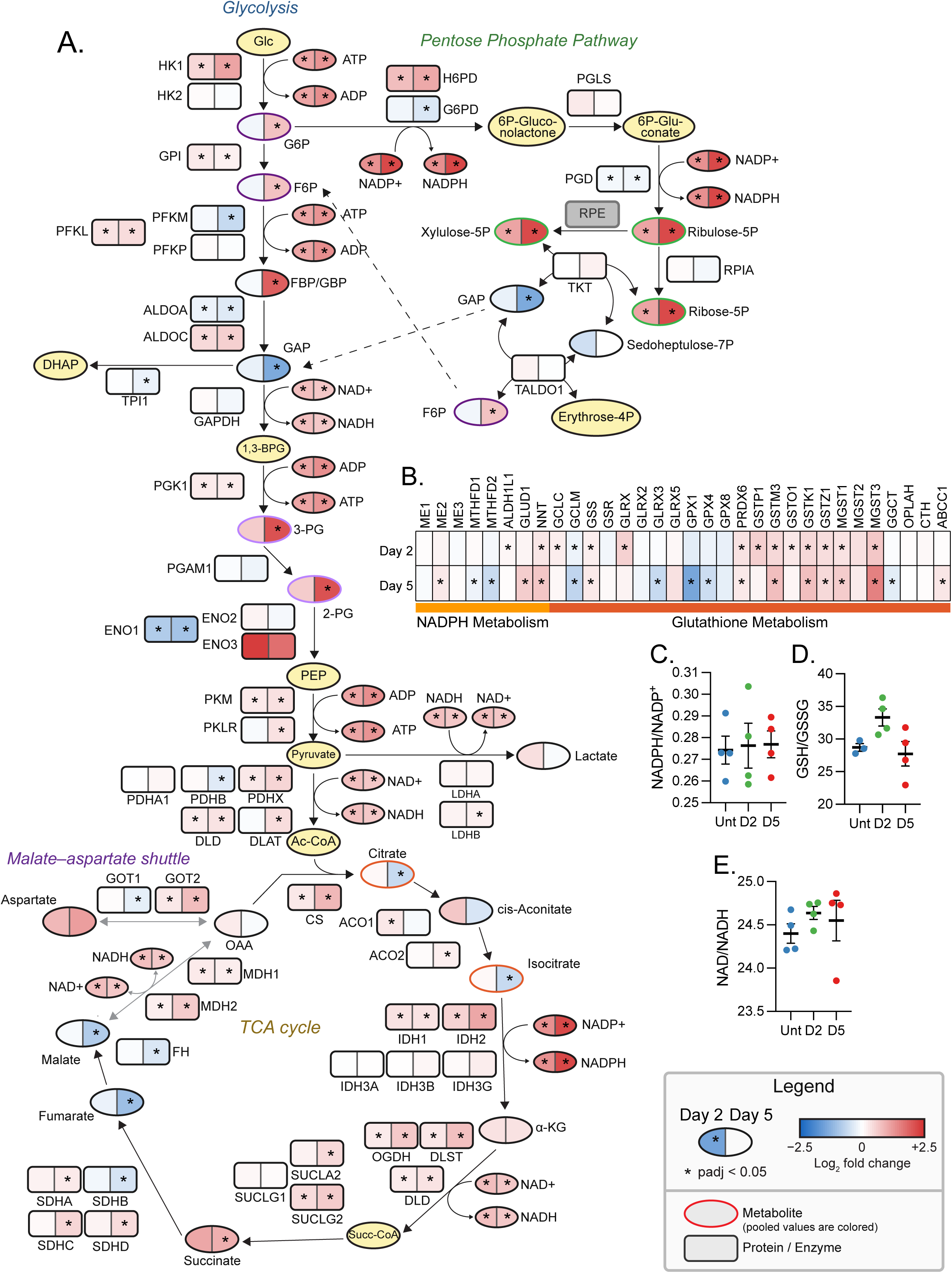
Heightened TCA cycle and antioxidant responses accompany oxphos elevation in radio-residual cells. (A) Glycolysis, pentose phosphate pathway, TCA cycle, and malate-aspartate shuttle pathway maps combining proteomic (rectangles) and metabolomic (ovals) profiling data. Color scale represents Log_2_ fold change of day 2 vs untreated (left side) and day 5 vs untreated (right side) samples and *padj<0.05 by BH corrected moderated t-test, n=4 (B) Heatmap displaying Log_2_ fold change of day 2 vs untreated and day 5 vs untreated proteins involved in NADPH and GSH metabolism *padj<0.05 by BH corrected t-test (C) NADP^+^/NADPH ratio, (D) GSH/GSSG ratio, and (E) NAD^+^/NADH ratio in untreated and day 2 and 5 post 8 Gy IR MDA-MB-231 cells quantified by mass spectrometry.

Metabolomics analyses uncovered substantial increases in redox metabolites (**Figure S5C,F).** While both NAD^+^ and NADH, NADP^+^ and NADPH, and GSSG and GSH were significantly increased, their ratios remained stable (**Figure 4C-E**). The maintained balance of these ratios points towards a heightened redox buffering capacity in oxphos-high radio-residual cells. Concordantly, we observed a proteomic upregulation in ‘NAD biogenesis and metabolism’ and ‘malate-aspartate shuttle’ pathways (**Figure S2**). Particularly, we observed significant elevation of nicotinamide nucleotide transhydrogenase (NNT), which generates NADPH and NAD^+^ from NADP^+^ and NADH, following IR (**Supplemental Table 5**). Similarly, elevation of oxidative pentose phosphate pathway (PPP) products ribulose-5-phosphate (ribulose-5P), xylulose-5-phosphate (xylulose-5P), and ribose-5-phosphate (ribose-5P), as well as an increase the NADPH producing enzyme hexose-6-phosphate dehydrogenase (H6PD) following IR suggested the PPP may be upregulated to mitigate oxidative stress (**Figure 4A**). We also observed increased GSH metabolism enzymes suggesting elevated ROS scavenging (**Figure 4B**). Together, our findings reveal that IR induced coordinated elevation of oxphos in tandem with metabolic rewiring, upregulating TCA cycle enzymes and metabolites as well as redox substrates and antioxidant capacity.

### IR elevates OPA1 protein isoforms associated with energy production and cristae formation

To gain a deeper understanding into mitochondrial rewiring in ‘radio-residual’ cells, we visualized the mitochondrial network in naïve and irradiated cells on days two, seven, and fifteen post-IR (**Figure 5A**). Quantification of mitochondrial morphology using MicroP^34^, revealed no substantial changes in mitochondrial elongation after IR (**Figure 5B**). Concordantly, immunoblotting revealed levels of mitochondrial fusion proteins MFN1 and MFN2 were unchanged throughout the study (**Figure 5C-D**). However, S616 phosphorylation of the mitochondrial fission protein DRP1, as well as isoform distribution of the mitochondrial shaping protein OPA1, were altered in residual cells (**Figure 5C-D**). The opposing contributions of OPA1 and DRP1 to mitochondrial fusion and fission, respectively, may explain the lack of substantial change in mitochondrial elongation or fragmentation after IR.

**Figure 5.**
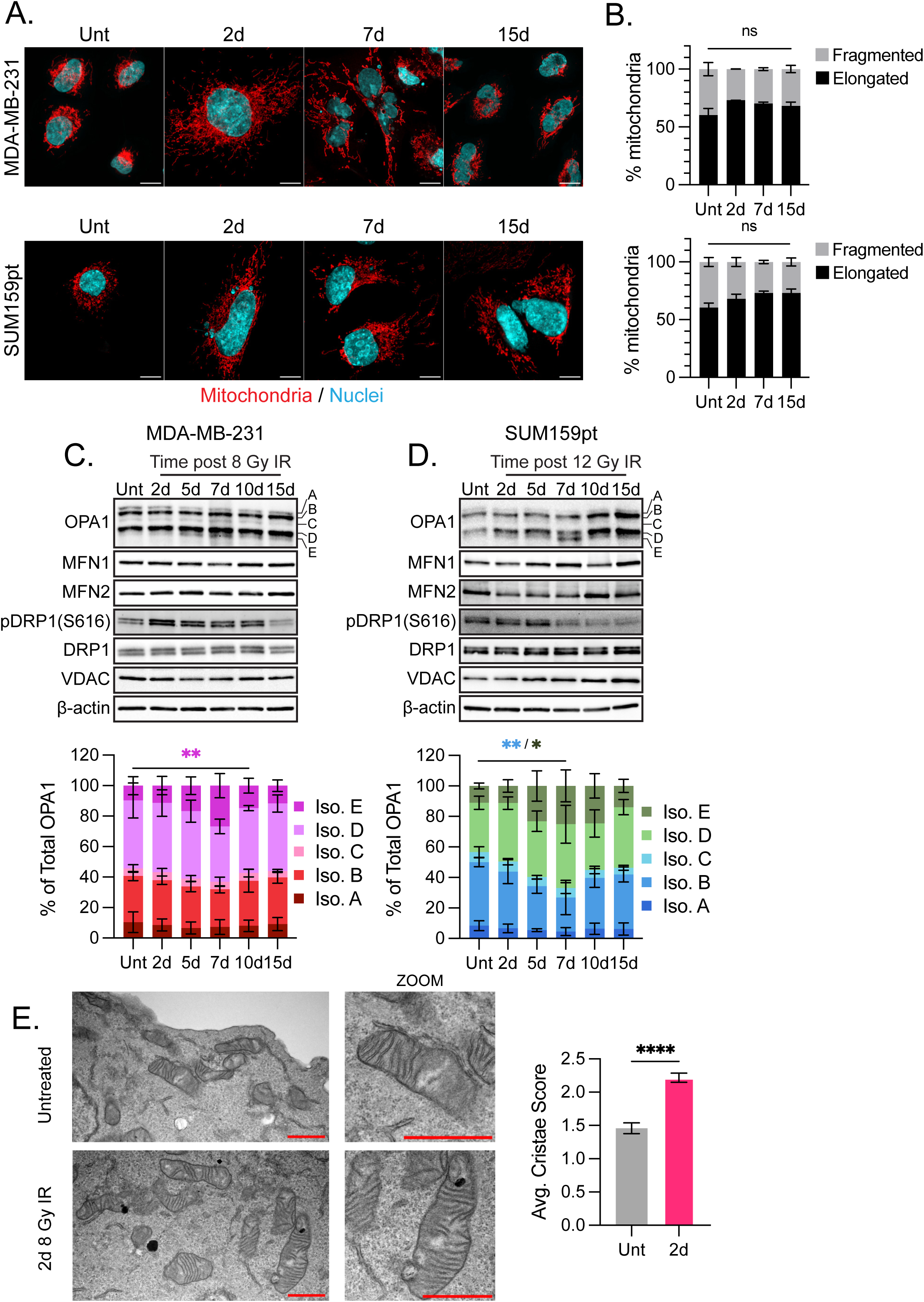
IR elevates OPA1 protein isoforms associated with energy production and cristae formation. (A) Representative fluorescent images of MitoTracker stained mitochondria (red) and DAPI-stained nuclei (blue) in untreated MDA-MB-231 and SUM159pt cells and at days 2, 7, and 15 post 8 Gy or 12 Gy IR, respectively. Scale is 10 µm. (B) Percentage of elongated and fragmented mitochondria after 8 Gy of IR in MDA-MB-231 cells and 12 Gy IR SUM159pt cells calculated using MicroP^34^. Data are represented as the mean ± SEM (n≥3, >15 cells per experiment). (C) Representative immunoblots (above) of mitochondrial dynamics proteins in 8 Gy IR MDA-MB-231 and (D) 12 Gy IR SUM159pt whole cell. Quantifications (below) of OPA1 isoform proportions represented as mean ± SEM. **P<0.01, *P<0.05 by Šídák’s multiple comparisons test comparing each isoform across each experimental group. Colored asterisks match isoforms in the legend. (E) Representative transmission electron micrographs of untreated and 2 days post 8 Gy IR MDA-MB-231 cells. Scale bar is 1 µm. Quantification of mitochondrial cristae scores for each experimental group are represented as mean score ± SEM of >249 mitochondria. ****P<0.0001 by Mann-Whitney U Test.

OPA1 produces five protein isoforms whose overlapping and non-overlapping functions are crucial for proper mitochondrial structure and function. Long OPA1 (L-OPA1) isoforms (A&B) have been shown to be the key isoforms driving mitochondrial fusion, while short OPA1 (S-OPA1) isoforms (C, D, &E) promote proper cristae formation and energy production^41^. In MDA-MB-231 cells L-OPA1 isoform levels were unchanged following IR, while S-OPA1 isoform E was significantly increased by day seven (**Figure 5C**). Interestingly, we observed a gradual elevation of both L-and S-OPA1 isoform levels post-IR in SUM159pt cells with a significant increase in the proportion of isoform E on day seven (**Figure 5D**). Given the role of S-OPA1 in regulating cristae structure, we sought to determine the impact of IR on cristae morphology using transmission electron microscopy (TEM). Mitochondria from MDA-MB-231 cells two days after IR had significantly increased cristae scores compared to mitochondria from untreated cells (**Figure 5E**), with a notably high proportion of mitochondria with more numerous, densely packed cristae. Together, these results demonstrate that irradiation induced mitochondrial inner membrane structural changes accompanying OPA1 isoform level changes and oxphos elevation in TNBC cells.

### Mitochondrial rewiring and regrowth of ‘radio-residual’ TNBC cells depend on OPA1

We sought to test the functional contribution of OPA1 in promoting the ‘radio-residual’ state using MDA-MB-231 cells in which OPA1 was knocked out (KO) by CRISPR/Cas9 (**Figure 6A**). OPA1 KO cells had reduced mitochondrial elongation (**Figure S6A-B**), oxphos (**Figure 6B-C**), mtDNA content (**Figure 6D**), and cristae scores (**Figure 6E**) compared to WT cells (**Figure 2&5**). Despite the reduction in OCR, OPA1 KO cells did not exhibit a compensatory increase in ECAR (**Figure S6C**). Importantly, the IR-induced increases in oxphos, mtDNA content, and cristae scores observed in WT cells (**Figure 2&5**), were ablated by OPA1 KO (**Figure 6B-E**). In fact, compared to nonirradiated, irradiated mitochondria in OPA1 KO cells had significantly fewer cristae with abnormal, onion-like morphology, which has been previously linked with poor mitochondrial functioning^42,43^. These findings suggest that OPA1 was required for IR-induced mitochondrial reprogramming.

**Figure 6.**
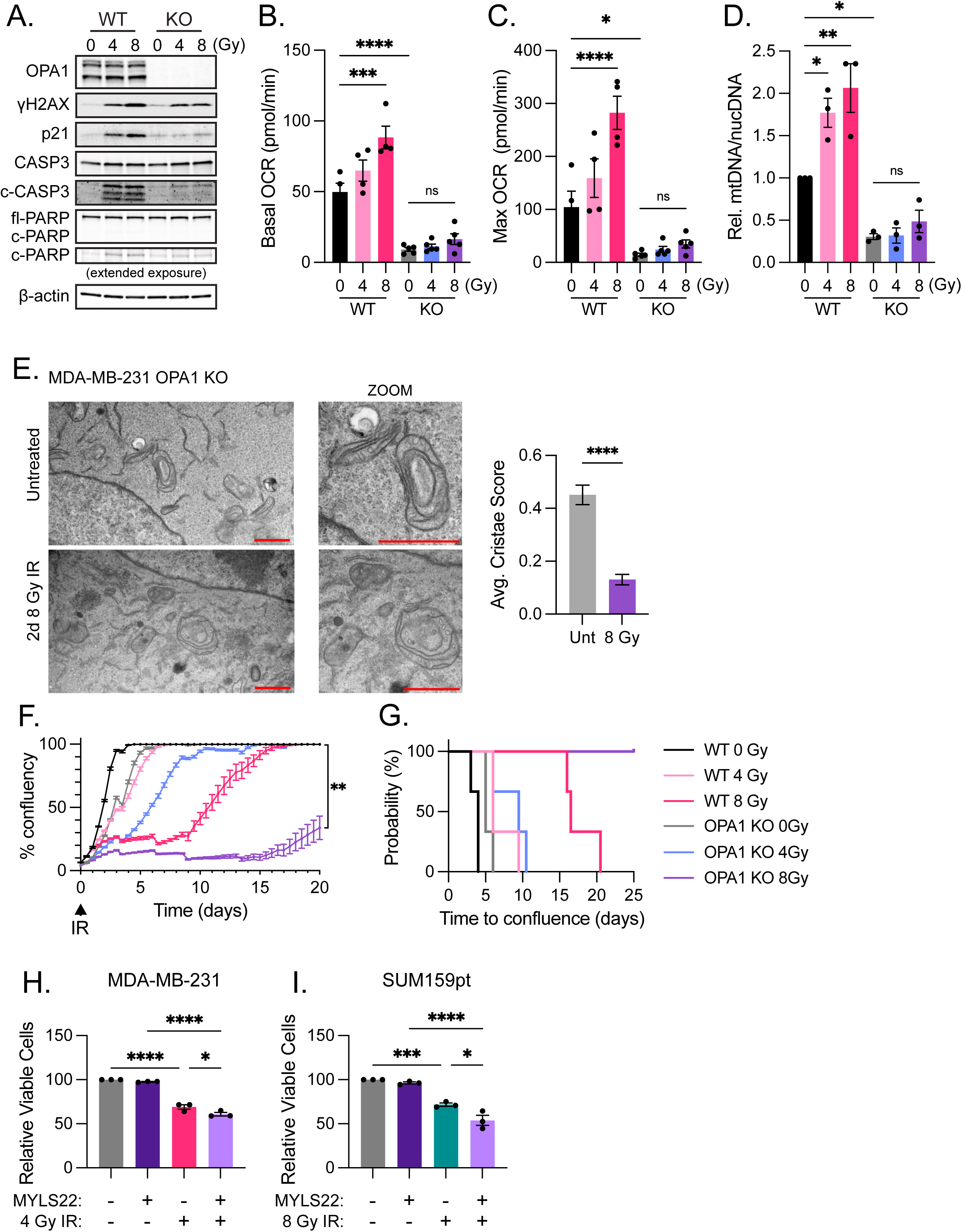
Mitochondrial rewiring and regrowth of radio-residual TNBC cells depend on OPA1. (A) Representative immunoblots in MDA-MB-231 WT and OPA1 KO cells treated with 0, 4, or 8 Gy collected at day 2 post-IR. (B) Basal OCR and (C) maximal OCR from Seahorse Mito Stress tests of MDA-MB-231 WT and KO cells treated with 0, 4, or 8 Gy collected at day 2 post-IR. (D) mtDNA content of MDA-MB-231 WT and KO cells **P<0.01, *P<0.05 by Šídák’s multiple comparisons test, n=3. (E) Representative transmission electron micrographs of untreated and 2 days post 8 Gy IR MDA-MB-231 OPA1 KO cells. Scale bar is 1 µm. Quantification of mitochondrial cristae scores for each experimental group are represented as mean score ± SEM of >326 mitochondria. ****P<0.0001 by Mann-Whitney U Test. (F) Representative Incucyte time lapse imaging of MDA-MB-231 WT and OPA1 KO cells treated with 0, 4, or 8 Gy IR. **P<0.01, by one sample t-test of day 20 confluence compared to 100, n=3 (G) Kaplan Meier plot generated from three independent Incucyte experiments as described in F. (H) Cell viability measurements on day 5 post-IR of MDA-MB-231 cells pre-treated with 50 µM MYLS22 for 24 hours then irradiated with 4 Gy as measured by Cell Titer-glo luminescence. (I) Cell viability measurements on day 5 post-IR of SUM159pt cells pre-treated with 50 µM MYLS22 for 24hr then irradiated with 8 Gy as measured by Cell Titer-glo luminescence. ****P<0.0001, *P<0.05 by Tukey’s multiple comparisons test, n=3.

OPA1 KO led to a pronounced delay in cell regrowth (**Figure 6F-G).** Despite OPA1 KO, IR still induced γH2AX accumulation, albeit to a lesser degree than in WT cells. KO cells exhibited reduced elevation of p21, cCASP3, and cPARP compared to WT cells, suggesting OPA1 KO cells may have altered cell cycle and cell death regulatory mechanisms (**Figure 6A**). As an orthogonal method to test the impact of OPA1 on IR sensitivity, we pre-treated MDA-MB-231 and SUM159pt cells with the small molecule OPA1 inhibitor, MYLS22^14^. MYLS22 treatment significantly reduced viability of irradiated cells, but not un-irradiated, SUM159pt cells, but only minutely did so in MDA-MB-231 cells (**Figure 6H-I**). Together, these findings provide evidence that OPA1 may functionally support IR-induced mitochondrial rewiring and survival of ‘radio-residual’ cells.

### Irradiation treatment rewires metabolism and mitochondrial processes in mouse mammary tumor models and human breast cancer cell lines

We next explored external datasets to test whether the mitochondrial adaptations we observed may be generalizable in TNBC. Although no electron microscopy datasets were available from unique TNBC models or patients, we identified a gene expression microarray dataset^35^ from cell lines derived from three unique p53-null syngeneic mouse mammary TNBC tumor lines^44–46^, that had been longitudinally profiled at several timepoints following treatment with 8 Gy of IR. We ran analogous mitochondrial gene expression analyses in those data side by side with our own proteomic dataset (**Figure S2**). Single-Sample Gene Set Enrichment Analysis (ssGSEA) of MitoCarta 3.0 pathways^22^ uncovered substantial similarities between the datasets. Specifically, luminal 2250L, basal like 2225L, and claudin-low T11 syngeneic models all had positive normalized enrichment scores (NES) in lipid, amino acid, and carbohydrate metabolism pathways beginning as early as 4 hours following IR and sustained through 48 hours (**Figure 7A; Supplemental Table 7**) matching our findings in MDA-MB-231 cells (**Figure S2**), which belong to the claudin-low subtype^47^. Mitochondrial central dogma pathways including mitochondrial translation, mtRNA metabolism, and mtDNA maintenance had positive NES acutely following IR in all models. However, NES declined with time in all three models, and the luminal model 2250L had significantly negative NES at 24 and 48 hours (**Figure 7A**). These findings align well with our MDA-MB-231 cell findings wherein mitochondrial central dogma related proteins were progressively reduced after IR (**Figure S2**). In summary, our findings suggest metabolic and mitochondrial reprogramming following IR, likely contributing to oxphos elevation and ‘radio-residual’ cell survival, may be a generalizable phenomenon in TNBCs.

**Figure 7.**
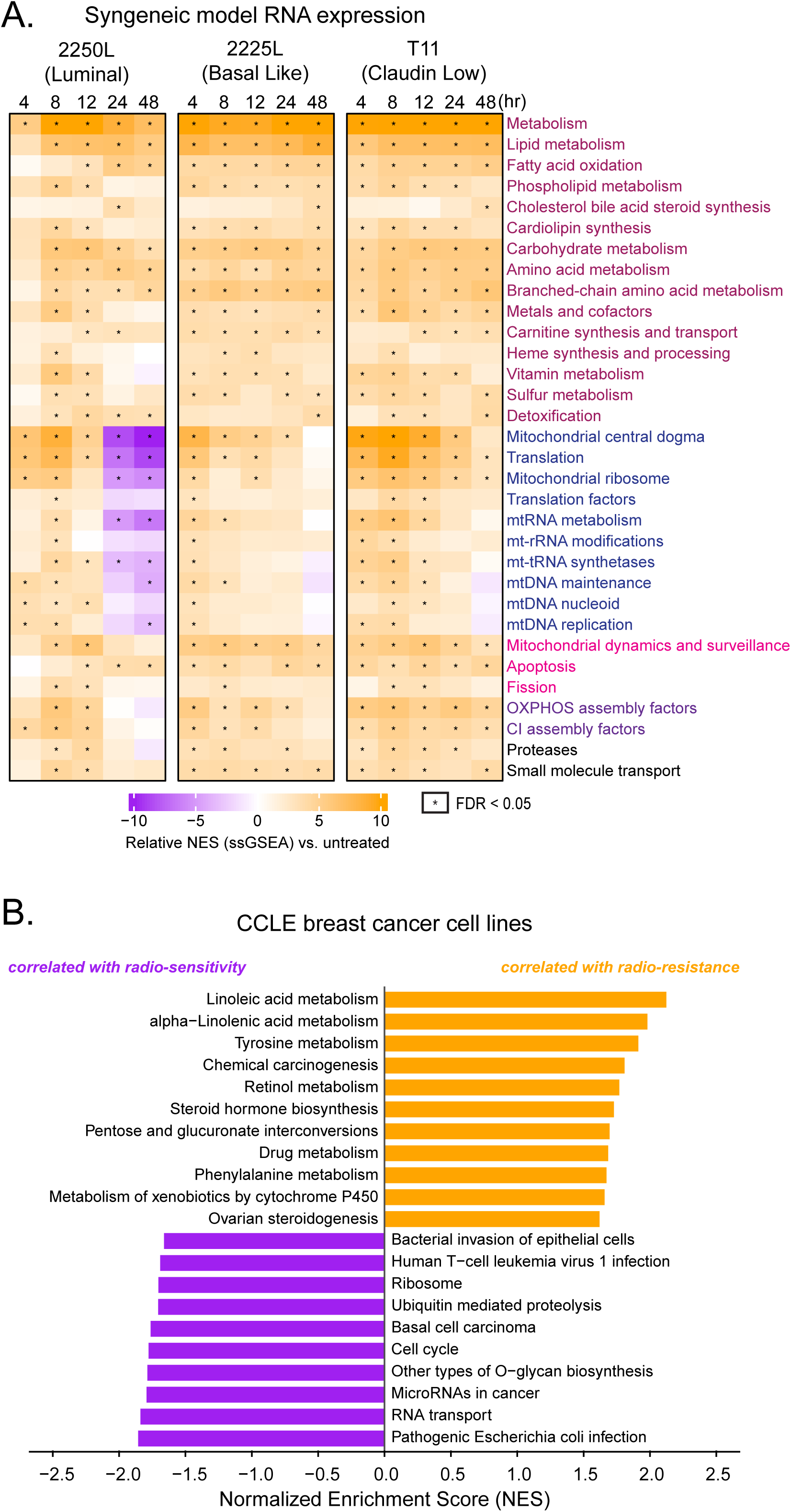
Metabolism and mitochondrial processes are altered in radio-residual mouse mammary tumors models and in radioresistant breast cancer cell lines. (A) Heatmap displaying ssGSEA NES scores of MitoCarta 3.0 pathways which had at least one significant sample in all syngeneic mouse models. (B) Bar plot showing normalized enrichment scores (NES) of KEGG pathways with FDR<0.05 from GSEA analysis of RNA expression correlated with radiation resistance of 28 breast cancer cell lines.

We also identified published IR response data for 533 cell lines cancer cell lines from the Characterized Cell Line Encyclopedia, including 28 breast cancer cell lines of which 12 were TNBC^48^. We then correlated publicly available RNA-seq gene expression data^49^ from the 28 breast cancer cell lines with IR response scores and performed GSEA of KEGG pathways. Cell lines in which expression of metabolic pathway genes were significantly more likely to have poor IR response and vice-versa for pathways associated with proliferation (*e.g.,* ribosome and cell cycle) (**Figure 7B**). In agreement with the metabolic pathways that became upregulated in radio-residual cells (**Figure S2**), the significantly correlation of innate metabolism-related gene expression with radioresistance in this external dataset affirms the potential importance of metabolic regulation in radiation-refractory TNBC.

## DISCUSSION

Radiotherapy is a mainstay of TNBC management, yet how tumor cells evade killing and ultimately relapse is poorly understood. This study provided new insights into the impact of plastic mitochondrial adaptations in TNBC cells that survive radiation exposure. We found that cells surviving IR undergo many metabolic adaptations to protect themselves from incurring cellular damage including slowing proliferation, upregulating oxphos, improving cristae density, increasing redox buffering, and mounting an antioxidant response.

Our study has inherent limitations. While our longitudinal model provides novel insights into molecular changes that occur when cells regrow, it should be noted that clinically, RT is given as fractionated doses rather than a single dose. While we can’t determine if small, fractionated doses equaling our IC_70_ doses would result in the same phenotypes we observed, it is important to note that lower doses resulted in similar mitochondrial adaptations as our IC_70_ dose (**Figure 6B-D**). Moreover, the mitochondrial rewiring we observed lasted at least one week, far exceeding the typical window between fractionated doses. Importantly, fractionated IR dosing of spheroid TNBC models recapitulated our results suggesting generalizability of the phenotype^37^. Another major challenge we faced was the lack of availability of human TNBC specimens or profiling datasets pre-and post-IR, due to the fact that irradiation usually occurs after surgical removal of the primary tumor, so there is typically no post-IR sample to be analyzed. In future studies, mitochondrial adaptations should be measured in additional TNBC models and, when possible, human biospecimens to ascertain the generalizability of our findings herein. It is also important to note that the effects of radiation on the tumor immune microenvironment are well established and are likely to be important for treatment efficacy. We were able to validate some of our findings in cell lines derived from immune-competent mouse models but future experiments dissecting mitochondrial rewiring in tumor and immune cells from *in vivo* TNBC mouse models will shed much needed light on this area.

Though our PDXO and *in vivo* data support mitochondrial adaptations following IR and our *in vitro* data implicate OPA1 as contributing to post-IR survival, we lack direct *in vivo* evidence of OPA1 inhibitors sensitizing TNBC to IR. The oxphos Complex I inhibitor IACS-10759 has been shown to increase radiosensitivity in NSCLC^50^ and intracranial sarcoma^50^ models supporting the efficacy of mitochondrial inhibitors to sensitize to IR. Another limitation of our study is the lack of direct evidence connecting OPA1-dependent elevation of cristae integrity to radio-sensitivity. Our data and existing literature suggest OPA1 is essential for cristae formation which is critical for ETC respirasome formation and oxphos efficiency^41,51–53^. Thus, we postulate OPA1 loss sensitizes cells to IR due to redox imbalance and elevated ROS production because of reduced respirasome formation. There is also evidence to suggest that IR induced oxphos supports survival simply by using up available oxygen that could otherwise form ROS^54,55^. Further investigations are needed to solidify this connection.

Oh and colleagues identified a radiation-induced gene signature that was predictive of pathological complete response (pCR) following chemotherapy, which provided us with a validation dataset^35^. Their data included RNA profiling of three p53-null syngeneic mouse cell lines representing, luminal, basal-like and claudin-low TNBC subtypes acutely following IR. Across all three models they observed similar changes to what we observed including upregulation of metabolic pathways and downregulation of proliferative pathways. Furthermore, our analysis of mitochondrial pathway changes in their data were very similar to our data. Lipid metabolism, carbohydrate metabolism, amino acid metabolism, mitochondrial dynamics, and ETC proteins are elevated in all models following IR. Mitochondrial central dogma pathways (*e.g.*, mtDNA replication, mtRNA metabolism, mito translation, etc.) show an initial elevation shortly after IR followed by a decline at different times in all three syngeneic models after IR (**Figure 7A**). Inhibition of mitochondrial transcription has been shown to sensitize cancer cells to IR especially those that are dependent on oxphos^56,57^. Proteomic analysis in our claudin low MDA-MB-231 cells align nicely with the RNA data claudin low model, T11, in that the mitochondrial ribosome pathway is unchanged 2 days post-IR. However, we see that by day 5 the mitochondrial ribosome pathway is decreased in MDA-MB-231 cells. Despite these findings, mtDNA-encoded ETC proteins were concomitantly elevated (**Figure 1H**), perhaps pointing towards a post-translational mechanism of protein level upregulation, such as increased protein stability. Indeed, proper cristae folding and ETC supercomplex formation have been demonstrated to promote the stability of ETC proteins within the complexes^52,58^, in alignment with the improved cristae morphology we observed in irradiated cells^59^ (**Figure 5E, 6E)**.

While targeting OPA1 and cristae formation has not been explored in the context of radio-resistance, it has been established that IR induces mitochondrial fission and elevation of fission proteins^60–63^. Indeed, we observe an increase in the fission pathway in our MDA-MB-231 proteomics as well as in our analysis of syngeneic mouse models RNA expression (**Figure 7A, S2**). Furthermore, we observe phosphorylated DRP (pDRP) is elevated following IR suggesting active fission, yet mitochondrial network morphology is not altered. This is likely due to simultaneous increases in OPA1 and fusion which leads to a net lack of change in mitochondrial elongation or fragmentation. These results suggest mitochondrial turnover rate is increased following IR to remove damaged mitochondria. The process of specifically recycling mitochondria is called mitophagy. Mitophagy, which is supported by DRP1, has been shown to be elevated in a variety of cancers^64–68^, including TNBC^62^, following IR. Furthermore, general autophagy can alter mitochondrial metabolism and has been shown to be activated by IR^69,70^. Corn *et al.* demonstrated that IR induced autophagy in TNBC fibroblasts led to metabolic adaptations that induced tumor cell growth^71^. Importantly, an autophagy inhibitor increased IR cell killing in a hepatocellular carcinoma model (HCC), providing proof of principle that autophagy inhibition can increase radiosensitivity^72^.

Prior studies have provided evidence that IR causes mitochondrial changes in tumor cells. While several studies have documented mitochondrial changes following IR in TNBC cells, contradictory findings have emerged ^37,56,73,74^. Our study sheds new light on the longitudinal, plastic nature of mitochondrial adaptations following IR and demonstrated the functional relevance thereof. Our work implicates OPA1 mediated cristae organization as a driver of radio-recurrence in TNBC and other metabolic pathways altered in response to IR. These findings merit further exploration beginning with OPA1 and extending to a variety of other mitochondrial and metabolism targeting strategies for their translational potential in TNBC.

## Acknowledgements

We are grateful to breast cancer patients who donated their biopsies for cell lines and PDX models. Ms. Janice Cowden and Dr. Amy Beumer, PhD provided patient research advocacy support for this work. Dr. Junegoo Lee, PhD, aided in radiation dose optimization for cell lines. Dr. Charles Perou, PhD provided consultation for mouse model microarray data analysis. Ariana C. Acevedo-Diaz, Luke Connell, MS, Denae Neill, and Alan Lopez Hernandez assisted with irradiation of cell cultures. Sahithi Puvvala measured cristae scores in electron micrographs. Dr. Mostafa Gaber, PhD, performed irradiation on tumors in mice. Dr. Alana Welm provided the PDXO model. Dr. Luca Scorrano developed the MYLS22 OPA1 inhibitor.

Core facilities at Baylor College of Medicine (BCM) and M.D. Anderson Cancer Center (MDA) supported this work. Fluorescence microscopy was conducted at the Optical Imaging & Vital Microscopy Core at BCM. Seahorse was conducted at the Mouse Metabolism and Phenotyping Core at BCM supported by NIH grants UM1HG006348, R01DK114356, and R01HL130249. STR cell line validation was conducted by the Cytogenetics and Cell Authentication Core at MDA. Immunohistochemistry was conducted at the Pathology Core and Lab in the Breast Center at BCM, which is supported by the Breast Center, and various other research grants including a NIH Specialized Programs of Research Excellence (SPORE) in Breast Cancer grant. ROS experiments were conducted at the Cytometry and Cell Sorting Core (CCSC) at BCM with funding from the CPRIT Core Facility Support Award (CPRIT-RP240432), the NIH (CA125123 and ODO36336) and the assistance of Joel M. Sederstrom. The OPA1 KO cell line was generated, and PDXO experiments were conducted at the Advanced Cell Engineering and 3D Models Core at BCM under the direction of Dr. Hugo Villanueva and Dr. Jun Xu. Proteomic and metabolomic profiling was conducted by BCM Mass Spectrometry Proteomics Core (RRID:SCR_027015) and Metabolomics Core. These facilities are supported by the Dan L. Duncan Comprehensive Cancer Center Award (P30 CA125123) and CPRIT Core Facility Awards (RP210227). The proteomics core is also supported by an Intellectual Developmental Disabilities Research Center Award (P50 HD103555) and a NIH High End Instrument Award (S10 OD026804, Orbitrap Exploris 480). We also acknowledge the joint participation of the Diana Helis Medical Research Foundation and the Adrienne Helis Malvin Medical Research Foundation and Baylor College of Medicine in support of the cores.

Proteomic and metabolomic profiling was conducted by BCM Mass Spectrometry Proteomics Core (RRID:SCR_027015) and Metabolomics Core. These facilities are supported by the Dan L. Duncan Comprehensive Cancer Center Award (P30 CA125123) and CPRIT Core Facility Awards (RP210227). The proteomics core is also supported by an Intellectual Developmental Disabilities Research Center Award (P50 HD103555) and a NIH High End Instrument Award (S10 OD026804, Orbitrap Exploris 480). We also acknowledge the joint participation of the Diana Helis Medical Research Foundation and the Adrienne Helis Malvin Medical Research Foundation and Baylor College of Medicine in support of the cores.

## AUTHOR CONTRIBUTIONS

S.W.W. and G.V.E were responsible for study conceptualization, oversight, experimentation, analyses, and manuscript preparation.

K.W. aided in initial project design and foundational data generation under the supervision of G.V.E.

A.G. conducted western blotting and animal studies under the supervision of G.V.E.

M.L.B. and M.J.B aided in mitochondrial morphology analyses under the supervision of G.V.E.

A.L. conducted computational analyses and OPA1 western blots under the supervision of G.V.E.

J.T.L. aided in computational analyses and figure construction.

A.A.G. aided in computational analyses under the supervision of G.V.E.

S.K. conducted western blotting and qPCR experiments under the supervision of G.V.E.

E.C. aided in radiation regimen design and treatments under the supervision of C.M.T. and E.S.

M.D.M. conducted transmission electron microscopy and aided in analyses.

S.L. conducted PDXO experiments under the supervision of H.V.

H.V. aided in PDXO experimental design and execution.

C.M.T. aided in initial project conceptualization and design.

R.P. provided project consultation and aided in animal study design.

