## Supplemental Tables and Figures for "Mitochondrial rewiring supports survival of triple negative breast cancer cells after ionizing radiation"

Supplementary Table 1

| Target | Source | Vendor | Cat. # | Application | Dilution |
|---|---|---|---|---|---|
| Ki67 | Mouse | Dako | 7240 | IHC | 1:200 |
| Cleaved Caspase 3 (CC3) | Rabbit | Cell Signaling Technology | 9661S | IHC | 1:50 |
| Phospho Histone H3 (pHH3) | Rabbit | Upstate | 06-570 | IHC | 1:250 |
| p21 | Mouse | Zymed | 18-0401 | IHC | 1:50 |
| Phospho Histone H2A.X (Ser139) (gH2AX) | Rabbit | Cell Signaling Technology | 2577S | WB | 1:1000 |
| p21 | Rabbit | Cell Signaling Technology | 2947S | WB | 1:1000 |
| Cleaved Caspase 3 (CC3) | Rabbit | Cell Signaling Technology | 9664S | WB | 1:1000 |
| Caspase 3 (CASP3) | Rabbit | Cell Signaling Technology | 9662S | WB | 1:1000 |
| PARP | Rabbit | Cell Signaling Technology | 9542S | WB | 1:1000 |
| SOD2 | Rabbit | Cell Signaling Technology | 13141S | IHC and WB | 1:1000 |
| TOM70 | Rabbit | Thermo Scientific | 14528-1-AP | WB | 1:1000 |
| VDAC | Rabbit | Cell Signaling Technology | 4661S | WB | 1:1000 |
| OPA1 | Rabbit | Cell Signaling Technology | 80471S | WB | 1:1000 |
| DRP1 | Rabbit | Cell Signaling Technology | 8070S | WB | 1:1000 |
| Phospho DRP1 (Ser616) (pDRP1) | Rabbit | Cell Signaling Technology | 3455S | WB | 1:1000 |
| MFN2 | Rabbit | Cell Signaling Technology | 11925S | WB | 1:1000 |
| MFN1 | Rabbit | Cell Signaling Technology | 14739S | WB | 1:1000 |
| beta-actin (ACTB) | Mouse | Cell Signaling Technology | 3700S | WB | 1:2000 |
| beta-tubulin (TUBB) | Rabbit | Cell Signaling Technology | 2146S | WB | 1:1000 |
| Anti-Mouse IgG H&L HRP-linked | Goat | Abcam | ab205719 | WB | 1:10000 |
| Anti-Rabbit IgG H&L HRP-linked | Goat | Abcam | ab205715 | WB | 1:10000 |

Supplementary Table 2

| Gene | Application | Forward Primer (5'-3') | Reverse Primer (5'-3') |
|---|---|---|---|
| OPA1 | PCR | TCCGGGTTTTCGATACGTGTG | CACTCCCCAAAGCACGTAAG |
| ND1 | qPCR | ATGGCCAACCTCCTACTCCT | TAGATGTGGCGGGTTTTAGG |
| ND6 | qPCR | TGGGGTTAGCGATGGAGGTAGG | AATAGGATCCTCCCGAATCAAC |
| RGPD1 | qPCR | GTGGAGCCACTGAGAATGGT | GCATGCCTGGCTGATTTTAT |
| FUNDC2P2 | qPCR | TGAGTCAGTGGACCTTGCAG | CAGAATGGTTTGCAAGCTGA |

### SUPPLEMENTAL FIGURE LEGENDS

**Supplemental Figure 1. Additional cell line radiation model reveals temporal cellular and molecular changes.** (A) Relative viable MDA-MB-231 and (B) SUM159pt cells quantified after AO/PI staining 2 days after 0, 2, 4, 6, 8, 10, and 12 Gy IR. (n=4) (C) Colony forming assays of MDA-MB-231 and SUM159pt cells treated with 0, 2, 4, 8, and 12 Gy IR. **P<0.01 by extra sum of squares test, n≥4 (D) Brightfield images of SUM159pt cells acquired longitudinally after 12 Gy of IR. Scale bar is 200 μm. (E) Relative viable SUM159pt cells counted after AO/PI staining after 2, 5, 7, 10, and 15 days post 12 Gy IR. Counts were normalized to untreated cells collected at day 2 post-IR. ***P<0.001, *P<0.05 by one sample t-test compared to 100, n≥3. (F) Mean cell volumes calculated from mean diameters of viable cells. No significant change by ANOVA. (G) Representative immunoblots in SUM159pt whole cell lysates collected at days 2, 5, 7,10, and 15 post 12 Gy IR. (H) DCFDA fluorescence measured by flow cytometry in MDA-MB-231 cells after 24 hours of exposure to 4 and 8 Gy IR and in (I) SUM159pt cells after 24 hours of exposure to 8 and 12 Gy IR. Data represent relative median fluorescence intensity normalized to untreated controls for each experiment. **P<0.01, *P<0.05 by one sample t-test compared to 100, n≥3. (J) Relative viable regrown SUM159pt cells counted by AO/PI staining 2 and 5 days after 12 Gy IR. Counts were normalized to untreated cells collected at day 2 post-IR. ***P<0.001, **P<0.01 by one sample t-test compared to 100, n=3.

**Supplemental Figure 2. Radiation treatment rewires metabolism and mitochondrial processes in TNBC cell line.** Heatmap displaying KEGG pathways by gene set variation analysis (GSVA). Pathways were sorted by ANOVA and the top 25 most variable were displayed, n=4.

**Supplemental Figure 3. Oxidative phosphorylation is elevated in radio-residual cells.** (A) Representative Seahorse Mito Stress test graphs of 8 Gy IR MDA-MB-231 and (B) 12 Gy IR SUM159pt cells measured at day 2, 7, and 15 post-IR. (C) Quantified spare respiratory capacity, (E) basal ECAR, (G) coupling efficiency, (I) non-mitochondrial OCR, and (K) proton leak of MDA-MB-231 cells post 8 Gy IR. (D) Quantified spare respiratory capacity, (F) basal ECAR, (H) coupling efficiency, (J) non-mitochondrial OCR, and (L) proton leak of SUM159pt cells post 12 Gy IR. ****$P<0.0001$, ***$P<0.001$, **$P<0.01$, *$P<0.05$ by Dunnet's multiple comparisons test following one-way ANOVA, $n\geq4$.

**Supplemental Figure 4. Superoxide dismutase 2 is elevated in radio-residual cells.** (A) Representative immunoblots of SOD2 and TOM70 proteins in 8 Gy IR-treated MDA-MB-231 and (B) 12 Gy IR-treated SUM159pt whole cell lysates.

**Supplemental Figure 5. Metabolic profiling panels reveal gradual increase in many metabolites following IR.** (A) PCA plots of mass spectrometry metabolomic analysis of central carbon metabolites, (B) carbohydrate metabolism metabolites, (C) and redox metabolites in MDA-MB-231 cells at days 2 and 5 post 8 Gy IR. (D) Heatmap displaying scaled $Log_2$ normalized abundance values for metabolites in the central carbon panel, (E) carbohydrate metabolism panel, and (F) redox panel.

**Supplemental Figure 6. OPA1 loss sensitizes TNBC cells to IR and impairs IR-induced phenotypes.** (A) Representative fluorescent images of MitoTracker stained mitochondria (red) and DAPI-stained nuclei (blue) in untreated and 8 Gy IR MDA-MB-231 OPA1 KO cells 2 days post-IR. Scale is 10 μm. (B) Percentage of elongated and fragmented mitochondria untreated and 8 Gy IR MDA-MB-231 OPA1 KO cells 2 days post-IR calculated using MicroP. Data are represented as the mean ± SEM ($n\geq3$, >15 cells per experiment). (C) Basal ECAR from Seahorse

Mito Stress tests of MDA-MB-231 WT and OPA1 KO cells treated with 0, 4, or 8 Gy collected at day 2 post-IR.

Figure S1.

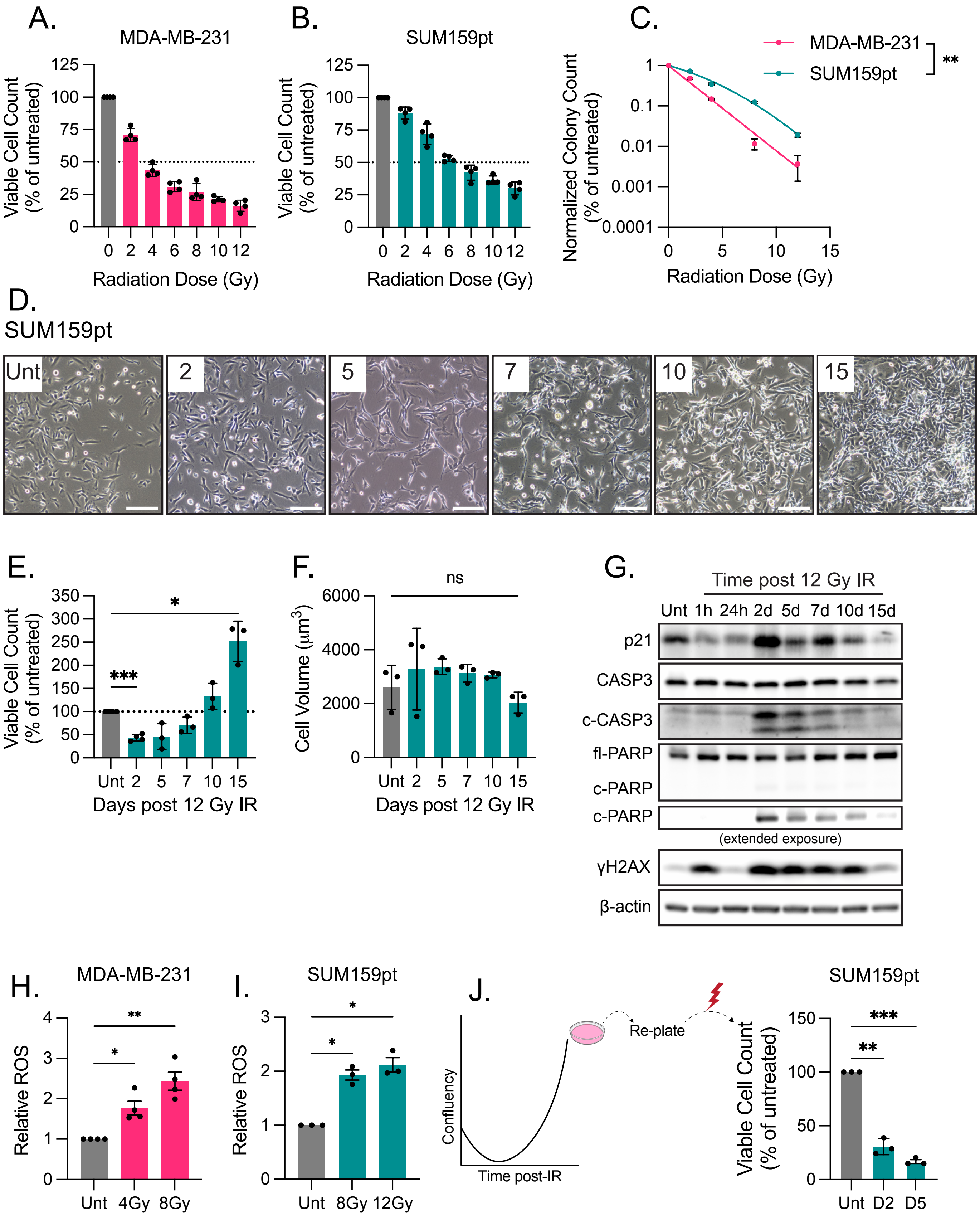
A.
MDA-MB-231
Viable Cell Count (% of untreated)
125
100
75
50
25
0
0 2 4 6 8 10 12
Radiation Dose (Gy)
B.
SUM159pt
Viable Cell Count (% of untreated)
0 2 4 6 8 10 12
Radiation Dose (Gy)
C.
MDA-MB-231
SUM159pt
**
Normalized Colony Count (% of untreated)
1
0.1
0.01
0.001
0.0001
0 5 10 15
Radiation Dose (Gy)
D.
SUM159pt
Unt
2
5
7
10
15
E.
Viable Cell Count (% of untreated)
350
300
250
200
150
100
50
0
*
***
Unt 2 5 7 10 15
Days post 12 Gy IR
F.
ns
Cell Volume (μm3)
6000
4000
2000
0
Unt 2 5 7 10 15
Days post 12 Gy IR
G.
Time post 12 Gy IR
Unt 1h 24h 2d 5d 7d 10d 15d
p21
CASP3
c-CASP3
fl-PARP
c-PARP
c-PARP
(extended exposure)
γH2AX
β-actin
H.
MDA-MB-231
Relative ROS
4
3
2
1
0
**
*
Unt 4Gy 8Gy
I.
SUM159pt
Relative ROS
3
2
1
0
*
*
Unt 8Gy 12Gy
J.
Confluency
Time post-IR
Re-plate
SUM159pt
Viable Cell Count (% of untreated)
150
100
50
0
***
**
Unt D2 D5

Figure S2.

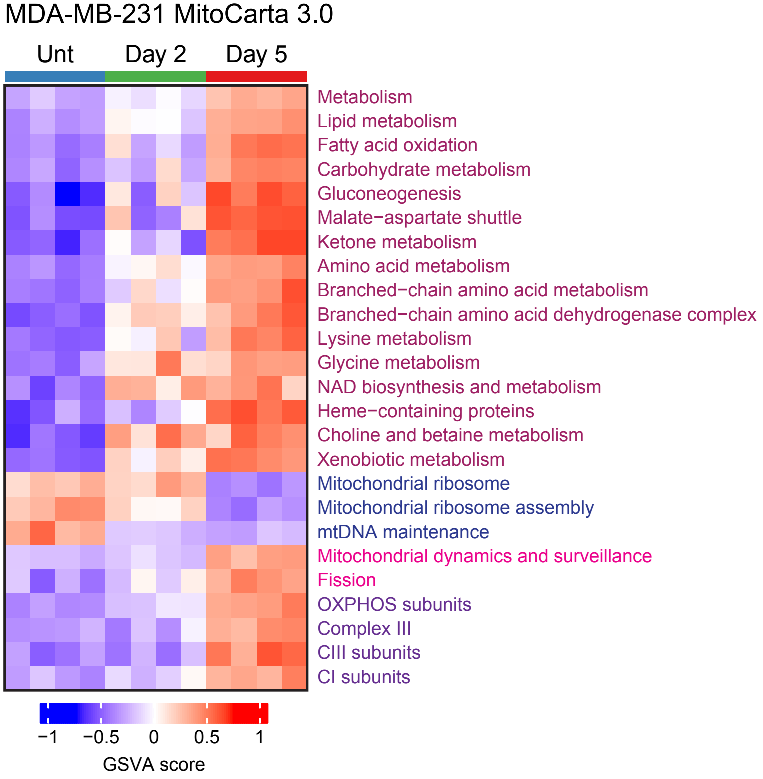
MDA-MB-231 MitoCarta 3.0
Unt
Day 2
Day 5
Metabolism
Lipid metabolism
Fatty acid oxidation
Carbohydrate metabolism
Gluconeogenesis
Malate−aspartate shuttle
Ketone metabolism
Amino acid metabolism
Branched−chain amino acid metabolism
Branched−chain amino acid dehydrogenase complex
Lysine metabolism
Glycine metabolism
NAD biosynthesis and metabolism
Heme−containing proteins
Choline and betaine metabolism
Xenobiotic metabolism
Mitochondrial ribosome
Mitochondrial ribosome assembly
mtDNA maintenance
Mitochondrial dynamics and surveillance
Fission
OXPHOS subunits
Complex III
CIII subunits
CI subunits
−1
−0.5
0
0.5
1
GSVA score

Figure S3.

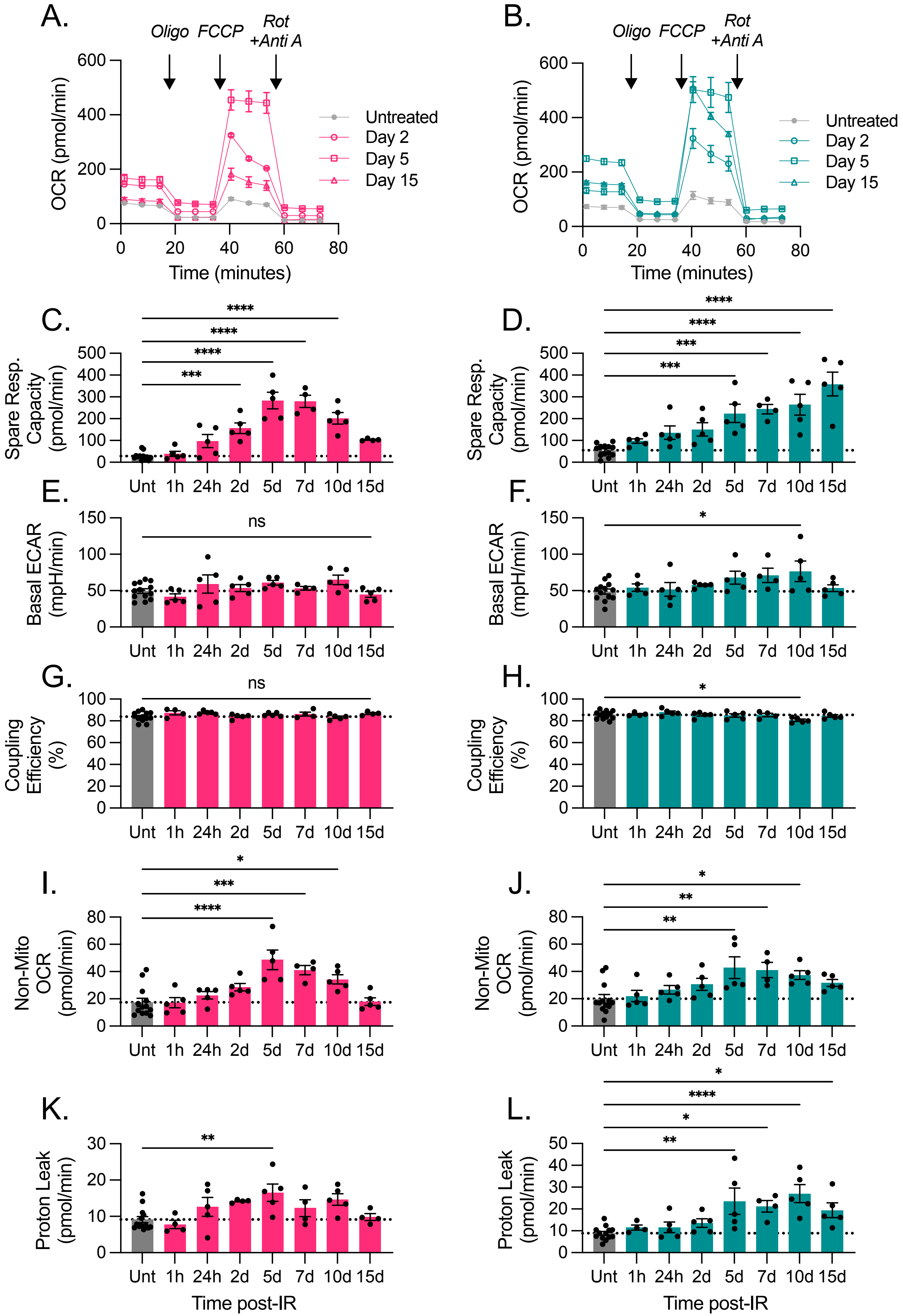

A.
Oligo
FCCP
Rot +Anti A
OCR (pmol/min)
Time (minutes)
Untreated
Day 2
Day 5
Day 15
B.
Oligo
FCCP
Rot +Anti A
OCR (pmol/min)
Time (minutes)
Untreated
Day 2
Day 5
Day 15
C.
Spare Resp. Capacity (pmol/min)
D.
Spare Resp. Capacity (pmol/min)
E.
Basal ECAR (mpH/min)
ns
F.
Basal ECAR (mpH/min)
G.
Coupling Efficiency (%)
ns
H.
Coupling Efficiency (%)
I.
Non-Mito OCR (pmol/min)
J.
Non-Mito OCR (pmol/min)
K.
Proton Leak (pmol/min)
L.
Proton Leak (pmol/min)
Unt 1h 24h 2d 5d 7d 10d 15d
Time post-IR
Time post-IR

Figure S4.

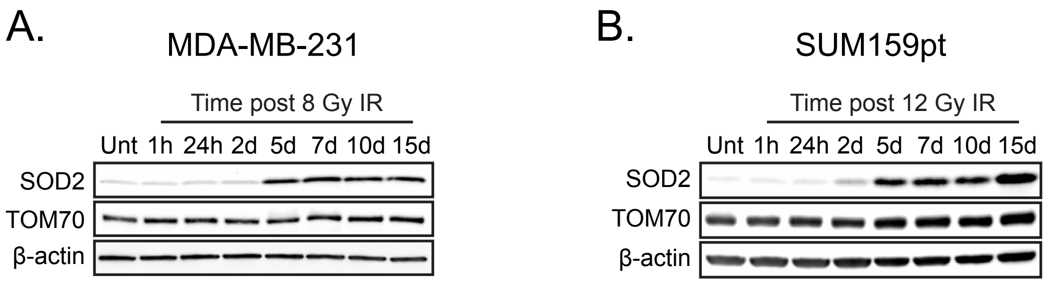
A.
MDA-MB-231
Time post 8 Gy IR
Unt 1h 24h 2d 5d 7d 10d 15d
SOD2
TOM70
β-actin
B.
SUM159pt
Time post 12 Gy IR
Unt 1h 24h 2d 5d 7d 10d 15d
SOD2
TOM70
β-actin

Figure S5.

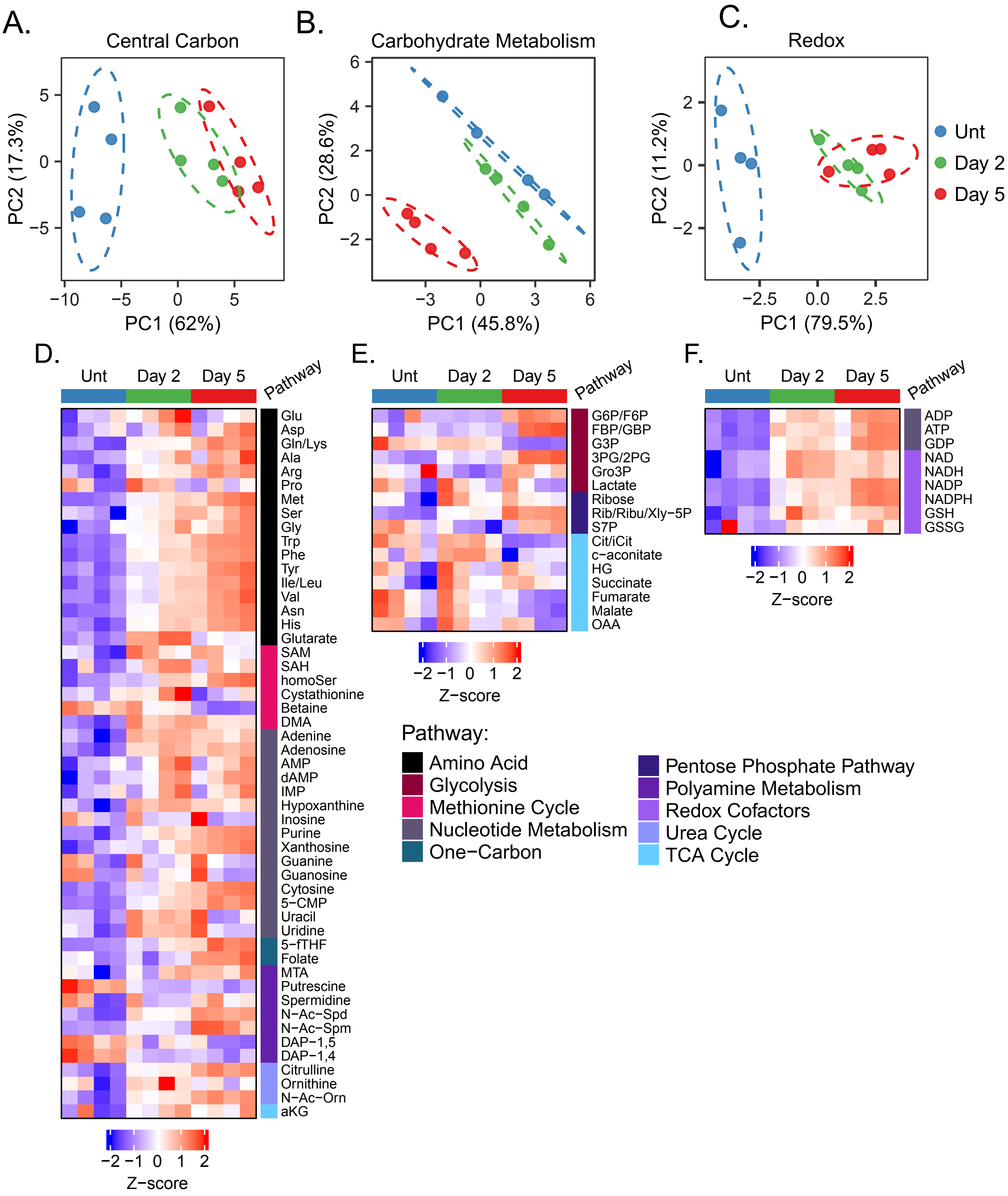
A.
Central Carbon
PC2 (17.3%)
PC1 (62%)
B.
Carbohydrate Metabolism
PC2 (28.6%)
PC1 (45.8%)
C.
Redox
PC2 (11.2%)
PC1 (79.5%)
Unt
Day 2
Day 5
D.
Unt
Day 2
Day 5
Pathway
Glu
Asp
Gln/Lys
Ala
Arg
Pro
Met
Ser
Gly
Trp
Phe
Tyr
Ile/Leu
Val
Asn
His
Glutarate
SAM
SAH
homoSer
Cystathionine
Betaine
DMA
Adenine
Adenosine
AMP
dAMP
IMP
Hypoxanthine
Inosine
Purine
Xanthosine
Guanine
Guanosine
Cytosine
5−CMP
Uracil
Uridine
5−fTHF
Folate
MTA
Putrescine
Spermidine
N−Ac−Spd
N−Ac−Spm
DAP−1,5
DAP−1,4
Citrulline
Ornithine
N−Ac−Orn
aKG
Z−score
E.
Unt
Day 2
Day 5
Pathway
G6P/F6P
FBP/GBP
G3P
3PG/2PG
Gro3P
Lactate
Ribose
Rib/Ribu/Xly−5P
S7P
Cit/iCit
c−aconitate
HG
Succinate
Fumarate
Malate
OAA
Z−score
F.
Unt
Day 2
Day 5
Pathway
ADP
ATP
GDP
NAD
NADH
NADP
NADPH
GSH
GSSG
Z−score
Pathway:
Amino Acid
Glycolysis
Methionine Cycle
Nucleotide Metabolism
One−Carbon
Pentose Phosphate Pathway
Polyamine Metabolism
Redox Cofactors
Urea Cycle
TCA Cycle

Figure S6.

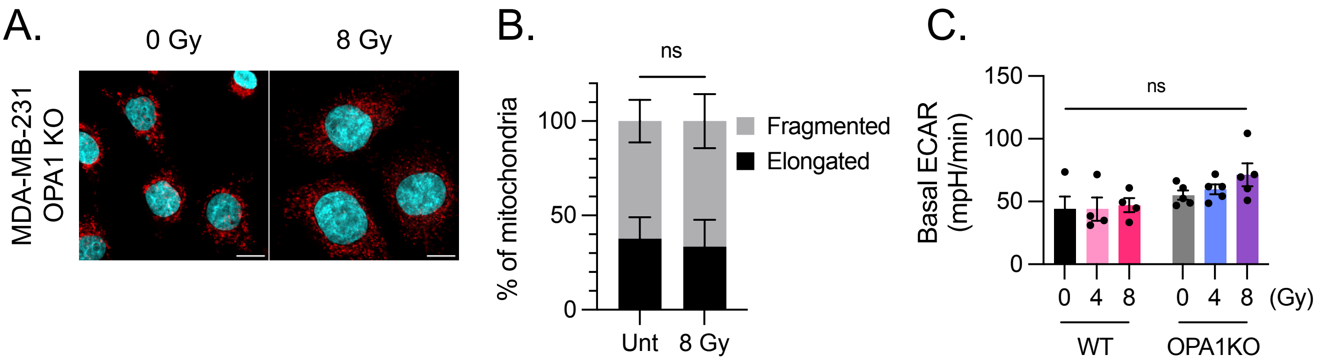
A.
0 Gy
8 Gy
MDA-MB-231
OPA1 KO
B.
ns
% of mitochondria
100
50
0
Fragmented
Elongated
Unt
8 Gy
C.
150
100
50
0
ns
Basal ECAR
(mpH/min)
0
4
8
0
4
8
(Gy)
WT
OPA1KO
